# Experienced meditators show greater forward travelling cortical alpha wave strengths

**DOI:** 10.1101/2024.06.09.598110

**Authors:** Neil W Bailey, Aron T Hill, Kate Godfrey, M. Prabhavi N. Perera, Jakob Hohwy, Andrew W. Corcoran, Nigel C. Rogasch, Bernadette M. Fitzgibbon, Paul B Fitzgerald

## Abstract

Mindfulness meditation involves training attention, commonly towards the current sensory experience, with an attitude of non-judgemental awareness. Theoretical perspectives suggest meditation alters the brain’s predictive processing mechanisms, increasing the synaptic gain and precision with which sensory information is processed, and reducing the generation or elaboration of higher-order beliefs. Recent research suggests that forwards and backwards travelling cortical alpha waves provide an indication of these predictive processing functions. Here, we used electroencephalography (EEG) to test whether the strength of forwards and backwards travelling cortical alpha waves differed between experienced meditators and a matched sample of non-meditators, both during an eyes-closed resting state (N = 97) and during a visual cognitive (Go/No-go) task (N = 126). Our results showed that meditators produced stronger forwards travelling cortical alpha waves compared to non-meditators, both while resting with their eyes closed and during task performance. Meditators also exhibited weaker backwards travelling cortical alpha wave strength while resting with their eyes closed. These results may be indicative of a neural mechanism underpinning enhanced attention associated with meditation practice, as well as a potential neural marker of the reductions in resting mind-wandering that are suggested to be associated with meditation practice. The results also support models of brain function that suggest attention modification can be achieved by mental training aimed at increased processing of sensory information, which might be indexed by greater strength of forwards travelling cortical alpha waves.

---

Mindfulness meditation involves training in focusing attention to the experience of the present moment, concurrent with an attitude of non-judgemental awareness (Kabat-Zinn, 2023). The practice of mindfulness meditation has been associated with improvements in well-being and improvements in some cognitive functions; improvements that are likely to be driven by changes in neural structure and activity (Bailey et al., 2020; Bailey, Freedman, et al., 2019; Boccia et al., 2015; Bowles et al., 2022; Chiesa et al., 2011; Ganesan et al., 2022). In particular, research has shown that meditation is associated with increases in grey matter volumes in specific regions, changes in metabolic activity in specific brain regions, increases in the power of neural oscillations, and altered neural activity during cognitive tasks (Bailey et al., 2023a; Fox et al., 2016; Fox et al., 2014; Lomas et al., 2015; McQueen et al., 2023; Osborn et al., 2022). However, our understanding of the neural mechanisms that underlie the effects of meditation is incomplete, with debate still underway about even the broader aspects of potential mechanistic explanations. For example, it is still uncertain how much the effects of meditation are due to increases in top-down attentional control processes, or due to alterations to bottom-up processing as a result of the mental training, or related to both of these mechanisms (Brandmeyer & Delorme, 2021; Chambers et al., 2009; Chiesa et al., 2013; Garland et al., 2009; Nakamura et al., 2021).

Recent theoretical perspectives have attempted to address this by explaining how meditation practice is likely to alter parameters of the predictive processing theory of brain function. The predictive processing framework suggests that because the brain only has indirect access to information about its environment (through sensory inputs to the brain), it acts as a hierarchical Bayesian inference model generator which constructs a predictive model of the brain’s environment and its place within that environment (Hohwy, 2020; Parr et al., 2022; Sprevak & Smith, 2023). Within the framework, the sensory perception, decision making, and motor functions of the brain are all suggested to be explainable by a single overarching driving force - the minimization of the long-term average prediction error (Friston, 2010; Hohwy, 2012). Minimization of prediction errors can be achieved through two routes. First, the brain can update prior beliefs in accordance with sensory evidence, making predictions more closely matched to incoming sensory input; this is perceptual inference (Friston, 2010; Hohwy, 2012; Sprevak & Smith, 2023). Second, the minimization of prediction errors can be achieved by using the brain’s motor control functions to direct the body’s musculature to orient the body such that the sensory organs selectively sample sensory input that conforms to prior beliefs; a process that is known as active inference (Parr et al., 2022). The minimization of this long-term average prediction error allows an organism to act to remain in preferred (or predicted) states for its own preservation and well-being (Pezzulo et al., 2022).

Within the predictive processing framework, prediction errors are weighted by their expected precision. To achieve this, increased synaptic gain is allocated to neural activity underlying the processing of prediction errors that are afforded more precision, such that the updating of prior beliefs is weighted in accordance with the degree of precision assigned to the prediction error. This precision weighting is thought to underpin and enable attentional functions (Feldman & Friston, 2010; Hohwy, 2012; Kok et al., 2012; Limanowski, 2022). Experimental research exploring neural markers of predictive processing indicates that (precision-weighted) prediction errors are passed up the neural hierarchy, with each level passing prediction errors via excitatory synaptic connections to the level directly above, and each level passing predictions to the level directly below via inhibitory synaptic connections that suppress predicted sensory evidence and allocate precision weighting (Parr & Friston, 2018; Sprevak & Smith, 2023). A simplification of the broader pattern of neural processes is that prediction error tied sensory inputs are predominantly processed in posterior regions of the brain, which are lower in the cortical hierarchy (Aitken et al., 2020; Gordon et al., 2019), while frontal brain regions that are higher in the cortical hierarchy are predominantly engaged in generating more abstract Bayesian predictions about expected sensory inputs (Hodson et al., 2023; Sprevak & Smith, 2023).

This characterisation of brain activity as a predictive processing system can be extended to explain other aspects of experience, including prospective forms of cognition such as decision-making and planning for the future. These processes can be explained as the brain’s attempt to minimize prediction errors between the brain’s preferred and predicted future sensations, as well as attempts to identify predictions that best match future events (Parr et al., 2022; Sprevak & Smith, 2023). Within this framework, top-down processes higher in the cortical hierarchy are suggested to relate to the experience of the “conceptual-self”, which processes abstractions including thoughts about past events and future events (which are counterfactuals that reflect mental time-travelling), while bottom-up sensory processes reflect the content of the more present-moment centred “experiencing-self” (Corcoran et al., 2020; Laukkonen & Slagter, 2021).

Within this predictive processing framework, the attention training aspect of meditation (which is commonly focused on bodily sensations) has been proposed to enhance the brain’s capacity to modulate precision-weighting towards hierarchically lower perceptual processes, by increasing synaptic gain for the weighting of prediction errors in inference (Laukkonen & Slagter, 2021; Lutz et al., 2019; Manjaly & Iglesias, 2020). The non-judgemental aspect of meditation has been suggested to reduce the production and elaboration of counterfactual predictive models (Deane et al., 2020; Lutz et al., 2019), reducing the magnitude and precision of the information flow down the neural hierarchy, from frontal regions to sensory regions (Laukkonen & Slagter, 2021). However, while these explanations fit elegantly with intuitions about how the different aspects of meditation training affect the brain to result in an altered subjective experience, very little empirical evidence is available to support these proposals. An opportunity to address this gap may be provided by the analysis of cortical travelling waves.

Theoretical research indicates that travelling oscillatory waves can be an emergent phenomenon within systems that have connections between spatially separated nodes, but with the connectivity restricted by distance (Ermentrout & Kleinfeld, 2001). Recent research suggests that in the human brain, cortical travelling alpha waves provide an indicator of the strength of information flow down and up the neural hierarchy and might index both attention and predictive processing functions (Lozano-Soldevilla & VanRullen, 2019; Muller et al., 2014; Pang et al., 2020). These cortical travelling waves show amplitude peaks and troughs that travel through the cortex periodically (Alamia & VanRullen, 2019; Pang et al., 2020). The interaction between two or more cortical travelling waves has been suggested to generate the spatially stationary temporal oscillations observable when measuring EEG activity at single electrodes, as the periodic overlap of two travelling wave peaks creates a temporary voltage peak at the spatial location of the overlap (Ingber & Nunez, 2011). For example, if the peaks of a forwards travelling wave originating in an occipital region and a backwards travelling wave originating from a frontal region overlap in a parietal region at a rate of 10 times per second, a 10Hz alpha oscillation might be detected in parietal electrodes. If an oscillation in the temporal domain within an occipital electrode is replicated in more frontal electrodes, but with a phase delay that is progressively increased as the distant from the occipital electrode increases, then the wave can be assumed to be travelling forwards (Alamia & VanRullen, 2024). In contrast, phase lags that progressively increase from frontal to occipital electrodes suggest a backwards travelling wave (Alamia & VanRullen, 2024). See Figure 1 for a depiction of a backwards travelling cortical alpha wave extracted from some of our eyes-closed resting data.

**Figure 1.**
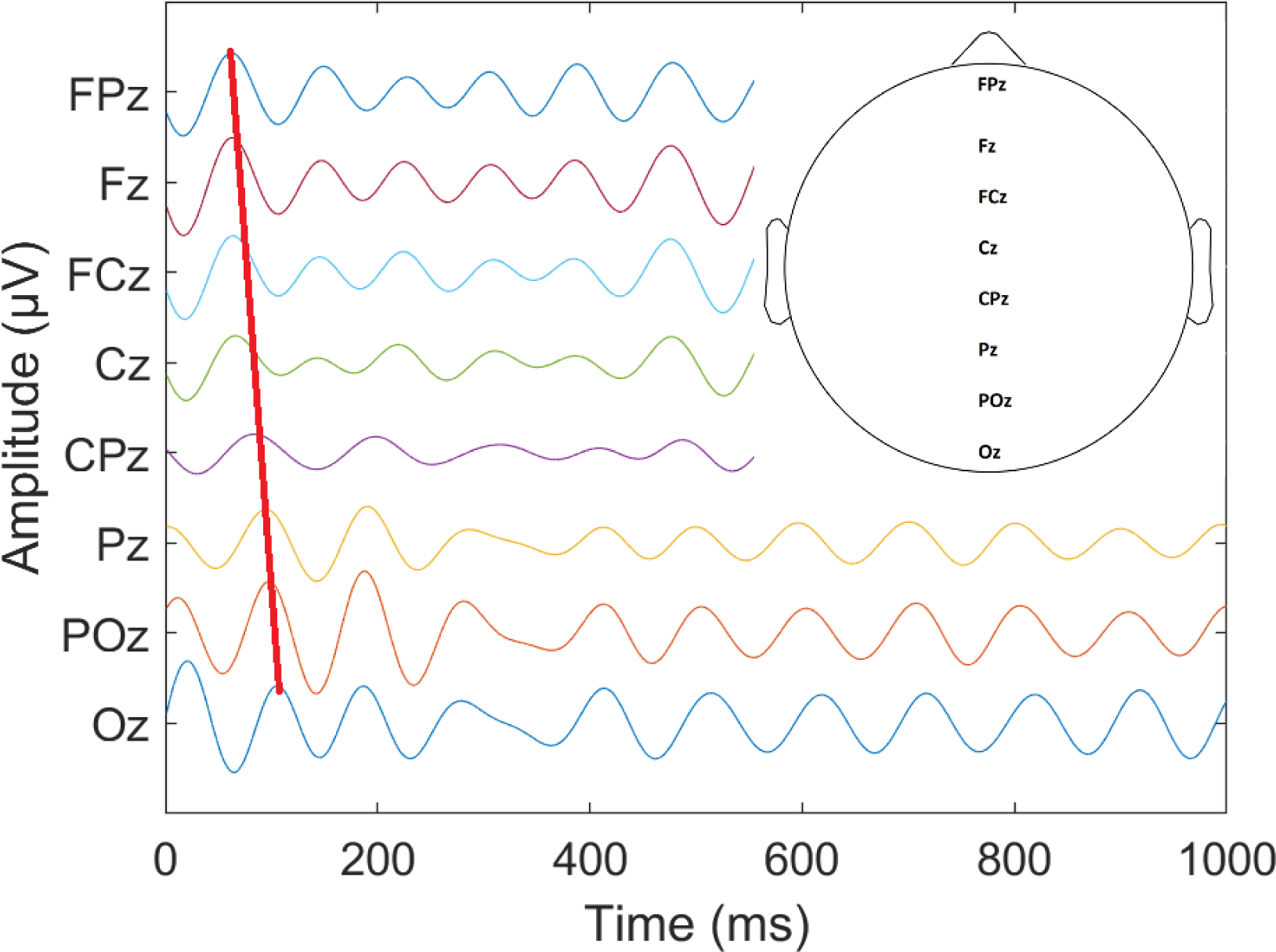
An example of a backwards travelling cortical alpha wave from real eyes-closed resting data bandpass filtered within the alpha band (8 to 13Hz). The red line highlights the oscillatory peaks, which are shifted progressively later in the more posterior electrodes. Right inset: The EEG electrodes included in the analysis.

Evidence from studies of visual attention tasks suggests that cortical travelling waves that originate in frontal regions and travel backwards provide an index of top-down predictions and are related to the direction of attention (Alamia & VanRullen, 2019; Pang et al., 2020). Modulation of these backwards travelling waves was found to be present regardless of whether visual stimuli were presented or not, suggesting the effect was driven by the intentional allocation of attention alone, without any processing of visual input required. This result indicates that top-down focused attention allocation is at least in part implemented by mechanisms that cause modulation of backwards travelling cortical alpha waves, which decrease when processing of an attended visual field is facilitated, and increase when processing of the non-attended visual field is inhibited (Alamia, Terral, et al., 2023). Backwards travelling waves have also been shown to be stronger during memory recall than during encoding (Mohan et al., 2024), concordant with the view that episodic memory recall is reliant on top-down mechanisms of gain-control over prediction error units (Barron et al., 2020). In contrast, forwards travelling waves that originate in posterior regions and travel forwards to frontal regions have been linked with bottom-up propagation of prediction errors driven by changes in sensory inputs (Alamia & VanRullen, 2019; Ermentrout & Kleinfeld, 2001; Pang et al., 2020). As such, forwards travelling waves are associated with the presentation of visual stimuli and are minimal in eyes-closed resting EEG recordings (Alamia & VanRullen, 2019; Pang et al., 2020). Forwards travelling waves have also been shown to be stronger during memory encoding than memory recall, perhaps suggesting that memory formation is reliant on feedforward neural interactions from sensory to frontal regions (Mohan et al., 2024).

Given that meditation has been suggested to reduce the generation or elaboration of top-down predictions, while increasing the precision weighting devoted to processing sensory information and bottom-up prediction errors, our current research tested whether experienced meditators show a lower strength of backwards waves and a greater strength of forwards waves compared to non-meditators. For our primary hypotheses, we expected: 1) that meditators would show stronger forwards waves during a visual Go/No-go task (which presented a rapid sequence of stimuli and required participants to respond to one type of stimulus while withholding their response to the other), reflecting increased processing of sensory information when such information is presented, and 2) that meditators would show weaker backwards wave strength, both during the visual Go/No-go task, and during eyes-closed resting, as a consequence of reduced top-down prediction.

Additionally, to further interrogate the functional relevance of travelling waves in memory function, we performed exploratory analyses of forwards and backwards wave strength from meditators and non-meditators during a modified Sternberg working memory task that separated memory set presentation, delay, and probe presentation periods. Within this task, the memory set presentation period required participants to attend to a string of eight letters, memorizing those letters. As such, we expected the memory set presentation period to elicit a greater strength of forward waves, as the visual information is transmitted to frontal regions, and a lower strength of backwards waves, so predictions do not interfere with visual processing. The delay period of this task was simply a blank screen, so would not be expected to generate forward waves, and might be expected to generate more backwards waves. The probe period required participants to recall whether the probe was present in the memory set, so we expected stronger backwards waves during the probe presentation (when participants attempted to discriminate whether the probe was in the memory set presented four seconds earlier) in alignment with previous research (Mohan et al., 2024).

Finally, we conducted an exploratory test of potential associations between travelling wave strengths and behavioural performance. In alignment with previous research (Mohan et al., 2024), we assumed that stronger forwards waves and weaker backwards waves during the memory set presentation would lead to better encoding of all memory stimuli. Additionally, in alignment with previous research (Mohan et al., 2024) we expected that stronger backwards waves during the probe period would be associated with higher performance when the probe was present in the memory set (a situation that requires matching of the remembered stimuli to the single probe stimuli). If supported, these hypotheses would provide additional support for the functional interpretation of travelling cortical alpha waves.

## Methods

### Participants

Both resting EEG data and task related EEG data were recorded across two separate studies of experienced meditators, which each examined potential differences between meditators and non-meditators in neural activity related to a range of cognitive tasks. The first study included a total of 34 experienced meditators and 36 healthy control non-meditators, and the second study included a total of 39 meditators and 36 healthy control non-meditators. Both studies included the recording of eyes-closed resting EEG data (Bailey et al., 2024; McQueen et al., 2023) and a Go/No-go task that presented Go and No-go trials with equal probability, with no more than three of each trial type presented consecutively (Bailey, Freedman, et al., 2019; Bailey et al., 2023b; Bailey, Raj, et al., 2019). However, due to time constraints within the recording sessions, not all participants provided both types of recordings, and some recordings were excluded due to artifact contamination. As such, the final analysed samples for each type of EEG recording differed in size, with 126 participants providing Go/No-go task related EEG recordings (69 meditators and 57 non-meditators), 97 participants providing eyes-closed resting recordings (50 meditators and 47 non-meditators), and 84 participants providing both types of recordings (47 meditators and 37 non-meditators). During the eyes-closed resting recordings, participants were instructed to simply rest with their eyes closed, not to meditate, but to just let their mind do as it pleased. This instruction was provided to enable characterisation of potential trait differences in resting EEG activity, rather than differences related to the meditation state. More details on these datasets can be found in Bailey et al. (2024) for the resting data and Bailey, Freedman, et al. (2019), Bailey, Raj, et al. (2019), and Bailey et al. (2023b) for the Go/No-go data. Additionally, 29 meditators and 29 non-meditators from the first study completed a modified Sternberg working memory task (Bailey et al., 2020). Within this task, all stimuli to be remembered were presented simultaneously, followed by a brief working memory delay period then a probe presentation period where participants responded as to whether the probe was contained within the memory set (Bailey et al., 2020).

**Table 1.**
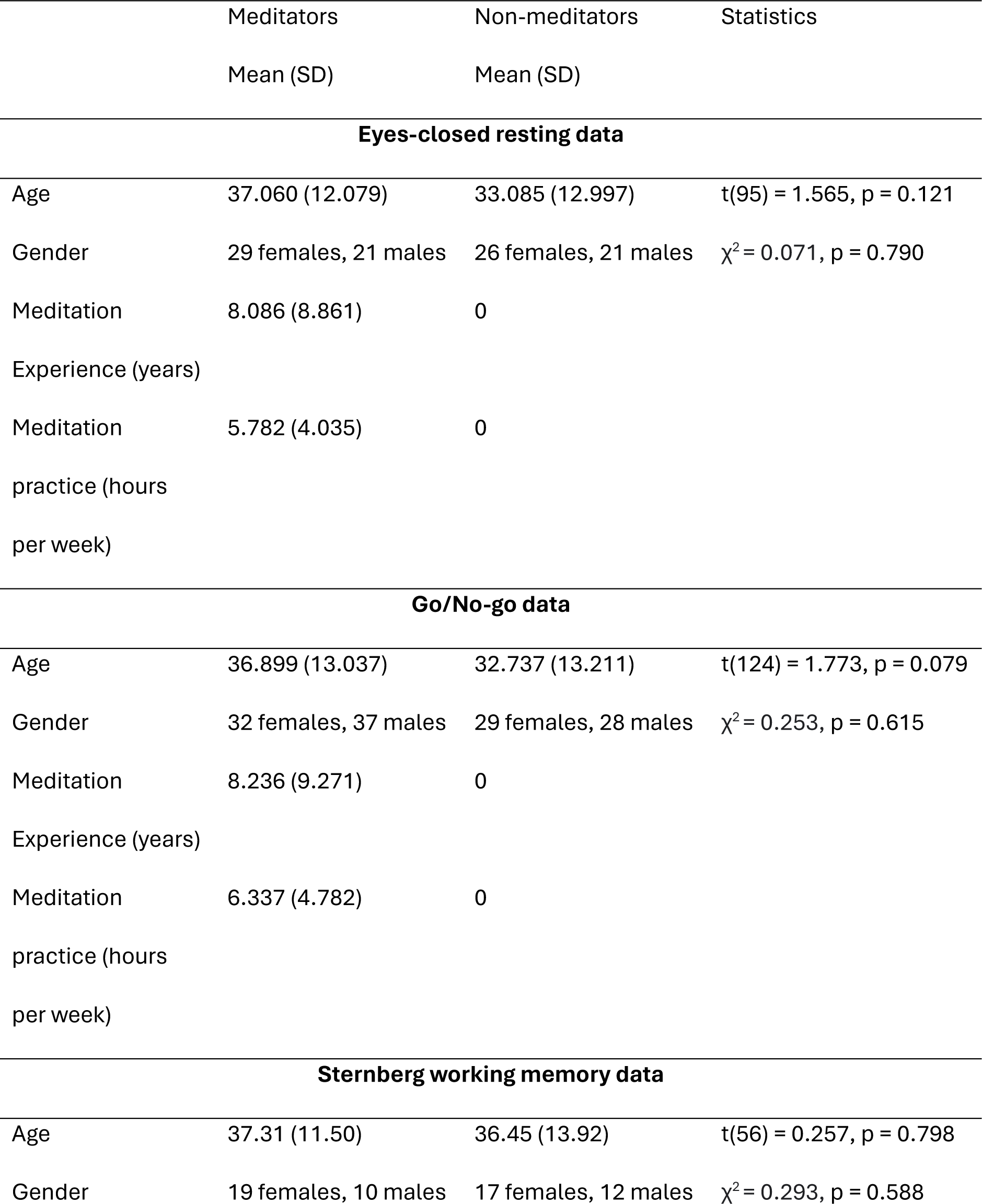
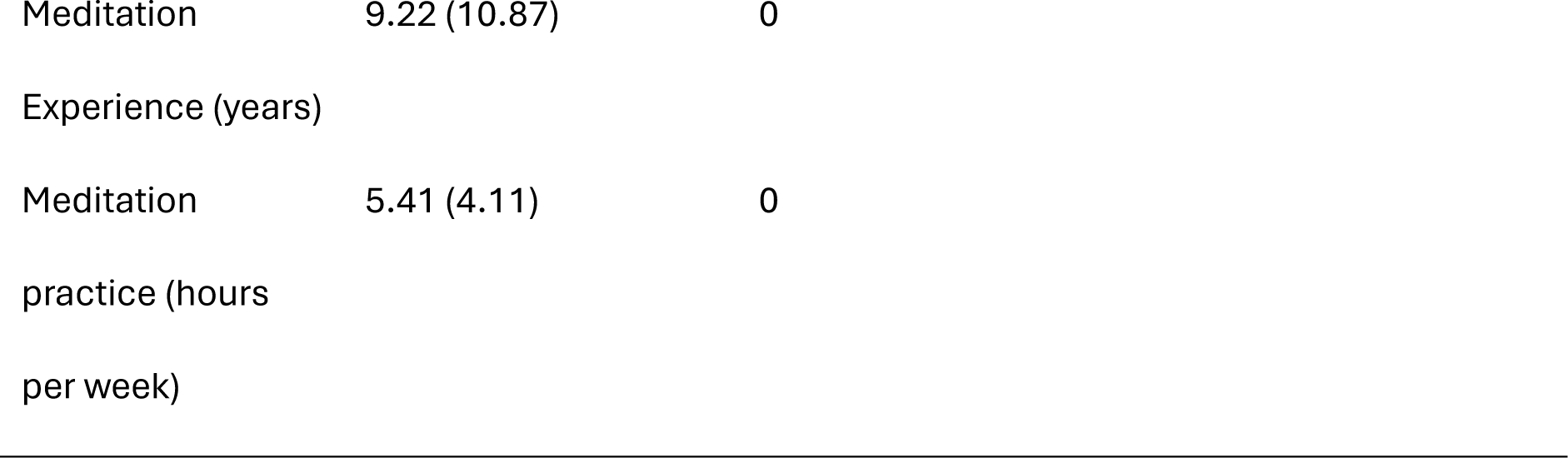
Demographic details for the different datasets.

|  | Meditators | Non-meditators | Statistics |
| --- | --- | --- | --- |
|  | Mean (SD) | Mean (SD) |  |
| **Eyes-closed resting data** |  |  |  |
| Age | 37.060 (12.079) | 33.085 (12.997) | $t(95) = 1.565, p = 0.121$ |
| Gender | 29 females, 21 males | 26 females, 21 males | $\chi^2 = 0.071, p = 0.790$ |
| Meditation | 8.086 (8.861) | 0 |  |
| Experience (years) |  |  |  |
| Meditation | 5.782 (4.035) | 0 |  |
| practice (hours<br>per week) |  |  |  |
| **Go/No-go data** |  |  |  |
| Age | 36.899 (13.037) | 32.737 (13.211) | $t(124) = 1.773, p = 0.079$ |
| Gender | 32 females, 37 males | 29 females, 28 males | $\chi^2 = 0.253, p = 0.615$ |
| Meditation | 8.236 (9.271) | 0 |  |
| Experience (years) |  |  |  |
| Meditation | 6.337 (4.782) | 0 |  |
| practice (hours<br>per week) |  |  |  |
| **Sternberg working memory data** |  |  |  |
| Age | 37.31 (11.50) | 36.45 (13.92) | $t(56) = 0.257, p = 0.798$ |
| Gender | 19 females, 10 males | 17 females, 12 males | $\chi^2 = 0.293, p = 0.588$ |

Table 1. Demographic details for the different datasets.
|  |  |  |
| --- | --- | --- |
| Meditation | 9.22 (10.87) | 0 |
| Experience (years) |  |  |
| Meditation | 5.41 (4.11) | 0 |
| practice (hours |  |  |
| per week) |  |  |

Both studies recruited participants through community advertising and advertising at meditation centres. Inclusion criteria for the meditators required a current mindfulness meditation practice with at least two hours per week of practice during the preceding two months, and a minimum of six months since they started practicing meditation. Meditation practices were required to be mindfulness-based, using the definition provided by Kabat-Zinn (2023): “paying attention in a particular way: on purpose, in the present moment, and nonjudgmentally”. Meditation practices were additionally required to have a practice component that involved focused attention on the breath of bodily sensations. Participants in the non-meditator group were required to have less than 2 hours of lifetime experience with any type of meditation. Participants who self-reported current or a history of mental or neurological illness or current psychoactive medication or recreational drug use were excluded. Participants were also excluded if they met criteria for any DSM-IV illness or scored in the moderate or higher range on the Beck Depression Inventory or Beck Anxiety Inventory. All participants were aged between 19 and 65 years of age and had normal or corrected to normal vision. All participants provided informed written consent prior to participation. The study was approved by the ethics committee of the Alfred Hospital and Monash University (approval number 194/14) and was conducted in accordance with the Declaration of Helsinki.

### EEG recordings

A Neuroscan 64-channel Ag/AgCl Quick-Cap was used to record EEG data using Neuroscan Acquire software and through a SynAmps2 amplifier (Compumedics, Melbourne, Australia). Online, electrodes were referenced to an electrode between Cz and CPz. Electrode impedances were reduced to below 5 kΩ prior to the start of each recording. EEG data were recorded at 1000Hz with an online bandpass filter of 0.05 to 200Hz. Data were pre-processed offline in MATLAB (The Mathworks, Natick, MA, 2023a) using EEGLAB (Delorme & Makeig, 2004), with data cleaning implemented using the default settings for the wICA_ICLabel cleaning approach of the RELAX pre-processing pipeline (Bailey, Biabani, et al., 2023; Bailey, Hill, et al., 2023). Briefly, this cleaning method first band-pass filtered the data from 1 to 80Hz and notch filtered the data from 47 to 53Hz using fourth order acausal Butterworth filters. RELAX then excluded bad electrodes then bad periods of the data using extreme outlier detection methods (including extreme outlier detections of: absolute voltages, voltage shifts, and probability distributions of values within each 1 second period). Data were then re-referenced to the robust average reference. Following these steps, independent component analysis (ICA) was performed to decompose the data using the PICARD algorithm (Ablin et al., 2018; Frank et al., 2022), and ICLabel was used to identify artifact components (Pion-Tonachini et al., 2019). The activations for each of these artifact components were then transformed using a stationary wavelet transform to characterize the dominant frequencies within each artifact component, which are assumed to reflect the artifact contribution to the signal (Castellanos & Makarov, 2006). The wavelet transformed versions of the artifact components were then subtracted from the raw artifact component with the intention to remove the artifact contribution to each component while preserving the neural activity (Bailey, Biabani, et al., 2023; Bailey, Hill, et al., 2023). The data were then reconstructed into the scalp space, providing data that were cleaned of artifacts in preparation for computation of the strength of travelling waves.

### Travelling wave computation

To determine the strength and direction of cortical travelling alpha travelling waves, we applied the method developed by Alamia and VanRullen (2019). Within each participant, 1 second epochs were extracted from each of the datasets. In alignment with the methods introduced by Alamia and VanRullen (2019), these epochs were extracted every 500ms within the resting data (so the epochs from the resting data contained 500ms overlaps with neighbouring epochs). For the Go/No-go data, epochs were time-locked to the stimulus presentation for all stimuli where participants provided a correct response (or non-response for No-go trials). A schematic of the tasks and the analysed periods can be viewed in Figure 2. Note that although the Go/No-go task presented stimuli every 900ms with a 50ms random jitter, we used 1 second epochs for consistency with previous travelling wave analyses and consistency with our resting-state analyses. This means that the travelling wave computation for each epoch was influenced primarily by activity following each time-locked stimulus, but also by up to 150ms of processing the following stimulus. However, since we did not separately analyse the Go and No-go trials, our results can be assumed to relate to the completion of the Go/No-go task in general, without being confounded by this inclusion of a portion of the neural activity in response to the following trial. For the Sternberg task, the 1 second epochs were extracted from the middle of the memory set presentation, delay period, and probe periods for trials where participants provided the correct response to the probe stimuli. This meant that within the probe presentation period, we measured activity starting 500ms after the probe was presented. This avoided activity related to visual processing, which reflects a deliberate decision implemented to focus analyses on processes reflecting participants’ attempts to match their recall of the memory set to the probe (processes which we expected would generate stronger backwards waves). Within each of these 1 second epochs, the EEG signal from the seven midline electrodes (Oz, POz, Pz, CPz, Cz, FCz, FPz) were extracted, providing a 2-dimensional matrix (electrode x time). A 2-dimensional fast Fourier transform (2D-FFT) was then applied to each electrode x time matrix within each epoch separately. The output matrix of this 2D-FFT provides both temporal frequencies and spatial frequencies, with spatial frequencies represented in the vertical axis of the matrix. Waves propagating in the forwards direction (from occipital to frontal electrodes) are represented in the upper quadrant of the matrix, while waves propagating in the backwards (from frontal to occipital electrodes) are represented in the lower quadrant (Alamia & VanRullen, 2019). Next, we extracted the maximum value within the alpha band (8 to 13Hz) from each quadrant within each epoch to measure the forwards and backwards travelling waves (Alamia & VanRullen, 2019). To ensure these travelling wave values reflected real signals that exceeded simple random patterns in the data, we performed the same travelling wave computations on a surrogate null version of the data. These surrogate null versions were obtained by shuffling the order of the electrodes in the electrode x time matrix separately within each epoch prior to the wave computation followed by computation of the 2D-FFT on the null data (Alamia & VanRullen, 2019). This shuffling of the electrode order destroyed the ability of the analysis to detect the spatial pattern of travelling waves provided by progressively increasing phase lags in more frontal electrodes (for forwards waves) or more posterior electrodes (for backwards waves), while still preserving the temporal oscillatory pattern, providing a distribution that was matched to the real data for oscillatory power but reflecting a null distribution for the spatial structure (Alamia & VanRullen, 2019). Finally, to ensure our statistical comparisons focused on the signal strength of the forwards and backwards travelling waves above the null, we divided the values from the real data within each epoch by the values within the surrogate data for the same epoch, then multiplied the result by 10*log10 and averaged these values across all epochs within each participant. This provided a value for each participant and condition that reflected the ratio of the strength by which the real forwards and backwards waves exceeded the surrogate forwards and backwards waves, with values on a log scale (providing values in decibel units [dB]) (Alamia & VanRullen, 2019).

**Figure 2.**
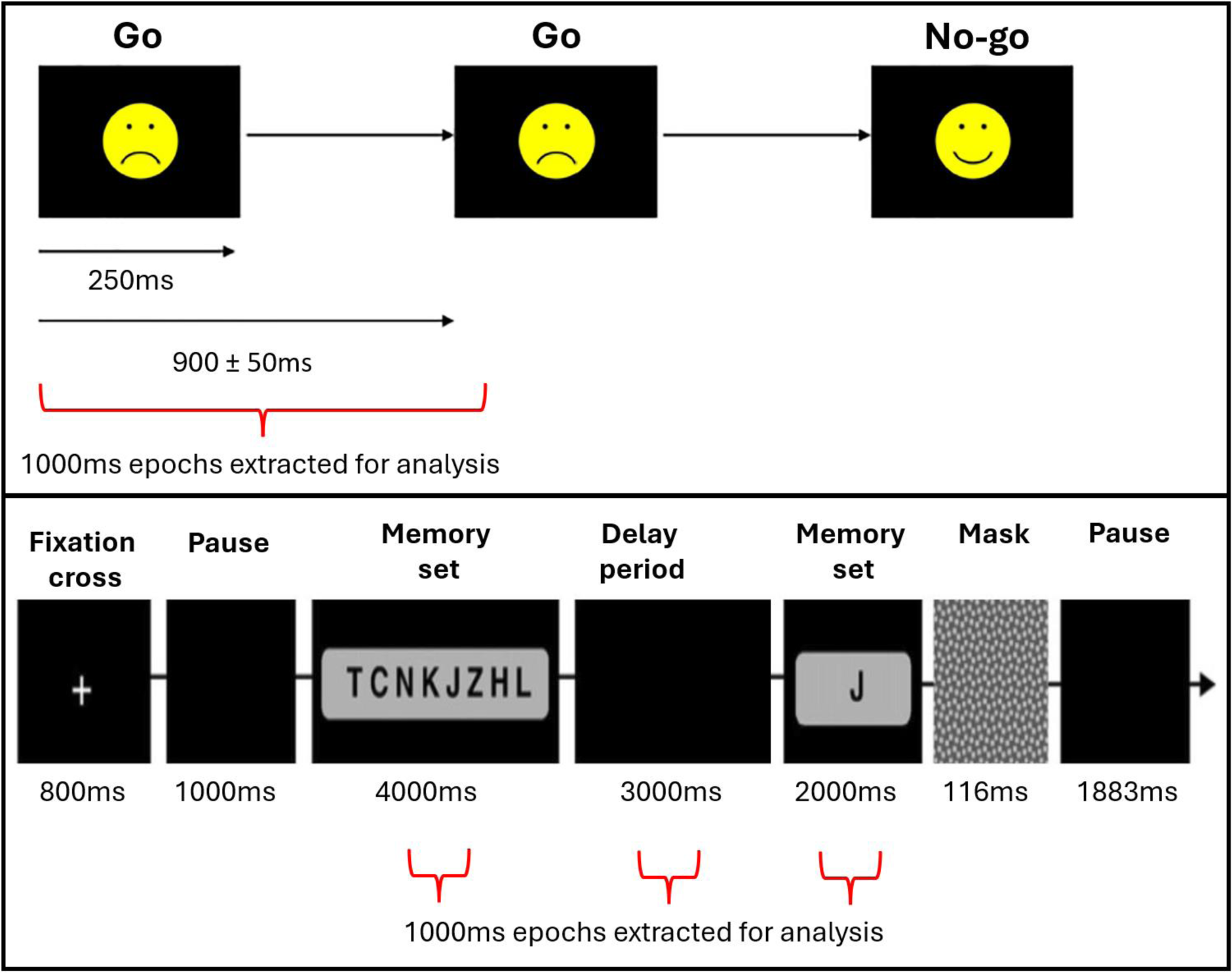
Schematic depictions of the tasks used to analyse forwards and backwards travelling wave strength. Top: the Go/No-go task presented emoticon style happy and sad faces in a 50:50 ratio, with stimulus response pairings switched between two randomly counterbalanced blocks so that all participants responded to an equal number of happy and sad faces and were required to withhold response to an equal number of happy and sad faces. Bottom: The modified Sternberg working memory task presented all memory set letters simultaneously and always contained a random selection of eight letters. The probe was contained within the memory set on 50% of the trials.

## Statistical Comparisons

Statistical analyses were performed using JASP 0.17.2.1 (Love et al., 2019). To control for the effect of outliers on our statistical analyses, prior to statistical analysis, we winsorized values that were more than three scaled median absolute deviations (MAD) from the median across all participants within the forwards and backwards waves for each condition separately. Winsorization was achieved by replacing the outlying value with the next most outlying value.

For our primary analysis, we conducted repeated measures ANOVAs separately within the resting and Go/No-go data to test for an interaction between group (meditators and non-meditators) and travelling wave direction (forwards and backwards), including all available participants in each ANOVA. To provide both a frequentist p-value and an indication of the strength of evidence for each analysis, we performed both frequentist ANOVAs and also Bayesian ANOVAs, and have reported the Bayes Factor (BF10) for each main effect and interaction effect compared to models stripped of that effect. Given that our expectation was for meditators to show stronger forwards waves and weaker backwards waves relative to controls, our primary focus was on the interaction between group and wave direction. As such, we controlled for multiple comparisons experiment-wise across these interactions for the eyes-closed resting and Go/No-go dataset using the false discovery rate method of Benjamini and Hochberg (1995), denoting corrected p-values as FDR-p. Additionally, we explored the drivers of significant interactions using post-hoc t-tests to make comparisons between the groups within specific conditions of interest. These post-hoc t-tests were also controlled for multiple comparisons using the false discovery rate.

In addition to these primary analyses, we performed several exploratory analyses to help with interpretation of the functional implications of the travelling waves. Firstly, within the sub-set of participants who provided both resting and Go/No-go data, we tested the correlation between forwards and backwards wave strength separately within each of these datasets across all participants. Matching the data across both recording types enabled us to assess potential differences in the strength of the relationship between the direction of the travelling waves between these recording conditions. Next, we tested for correlations between forwards and backwards wave strength and task accuracy in the Go/No-go dataset (with task accuracy measured by d-prime). After visual inspection of the data suggested a potentially interesting pattern, we also conducted exploratory repeated measures ANOVAs within the matched resting and Go/No-go dataset to test for interactions between the groups and EEG recording conditions (resting or Go/No-go) in the strength of the backwards waves.

Finally, to provide a deeper understanding of the potential functional relevance of the forwards and backwards waves, we explored the changing strengths of these waves during different periods of a sequential Sternberg working memory task. In particular, we tested wave strength within the memory set presentation period, delay period, and memory probe presentation period. To achieve this, we performed a repeated measures ANOVA with two within participant factors: wave direction (forwards or backwards) and working memory period (memory set presentation, delay period, and probe presentation period), and one between participant factor: Group. Finally, to explore whether these differences were functionally relevant, we inserted forwards and backwards wave strengths from each period of the working memory task as continuous predictors in a linear regression to predict working memory performance (measured by d-prime), with group as a between participant factor.

## Results

### Meditators show stronger forwards traveling waves and weaker backwards travelling waves during eyes-closed resting

Our analysis of cortical alpha travelling wave strength while participants were resting with their eyes closed showed that meditators exhibit stronger forwards travelling waves and weaker backwards travelling waves compared to non-meditators. This result suggests that meditation experience is associated with a greater strength of neural activity reflecting the bottom-up processing of sensory information and less generation of top-down neural activity when no task demands are present.

Specifically, within the eyes-closed resting data, the repeated measures ANOVA showed a significant interaction between group and wave direction: F(1,95) = 6.178, p = 0.015, FDR-p = 0.015, ηp² = 0.061, ηG² = 0.056, BFincl = 31.265 (Figure 3). Post-hoc t-tests showed that the interaction was driven by higher forwards wave strength within the meditator group (FDR-p = 0.022, Cohen’s d = 0.475, BF10 = 2.294), and lower backwards wave strength within the meditator group compared to the non-meditators (FDR-p = 0.022, Cohen’s d = 0.498, BF10 = 2.919). There was also a significant main effect of wave direction, with both groups showing higher values for the backwards wave than the forwards wave: F(1,95) = 50.966, p < 0.001, ηp² = 0.349, ηG² = 0.326, BFincl = 1.717*10^14^. The main effect of group was not significant: F(1,95) = 0.655, p = 0.420, ηp² = 0.007, ηG² < 0.001, BFincl = 0.178.

**Figure 3.**
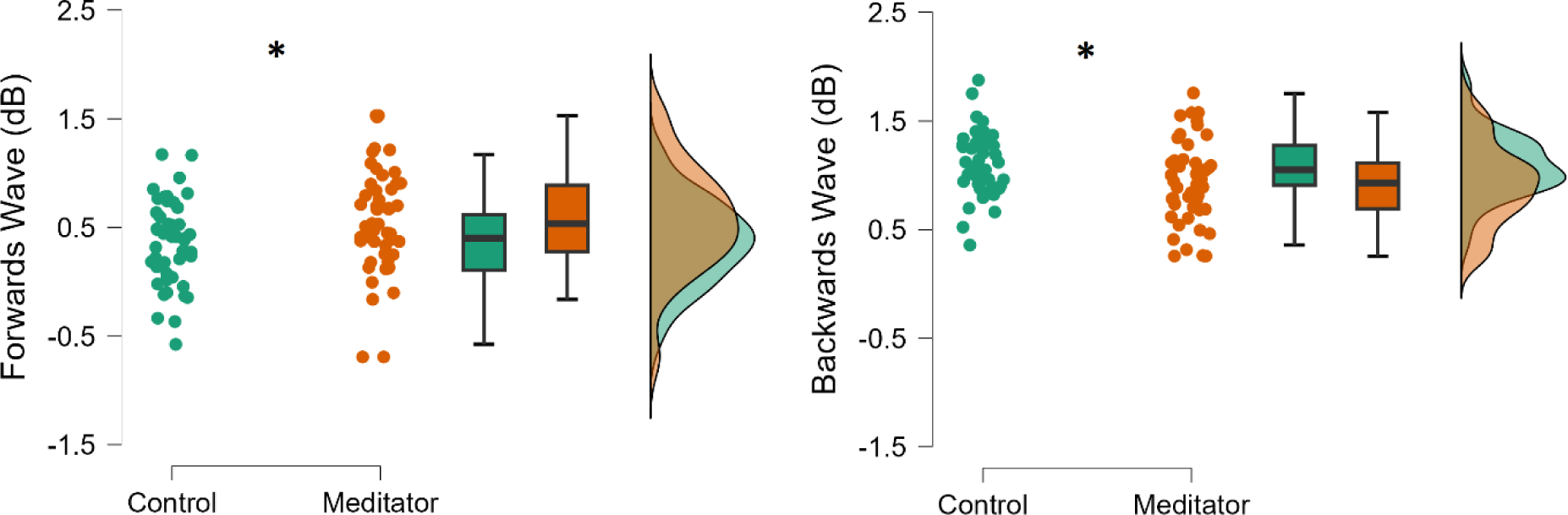
Forwards and backwards travelling cortical alpha wave strength within each group, measured from midline electrodes during the eyes-closed resting-state (values provided in decibels [dB]). Post-hoc testing showed that the interaction was driven by higher forwards wave strength within the meditator group (FDR-p = 0.022, Cohen’s d = 0.475, BF10 = 2.294), and lower backwards wave strength within the meditator group compared to the non-meditators (FDR-p = 0.022, Cohen’s d = 0.498, BF10 = 2.919). * FDR-p < 0.05.

Additionally, a strong negative correlation was present between the forwards and backwards wave strength in the resting data (r = -0.848, 95% CI = -0.774 to -0.899, p < 0.001, BF10 = 1.621*10^21^, see Figure 4). Note that we restricted this correlation to only the participants who provided data for both the eyes-closed resting recordings and the Go/No-go task to enable direct comparison of the correlation strength between the two recording types (N = 84). However, the full eyes-closed resting dataset showed an almost identical effect (r = -0.844, 95% CI = -0.776 to -0.893, p < 0.001, BF10 = 2.176*10^24^).

**Figure 4.**
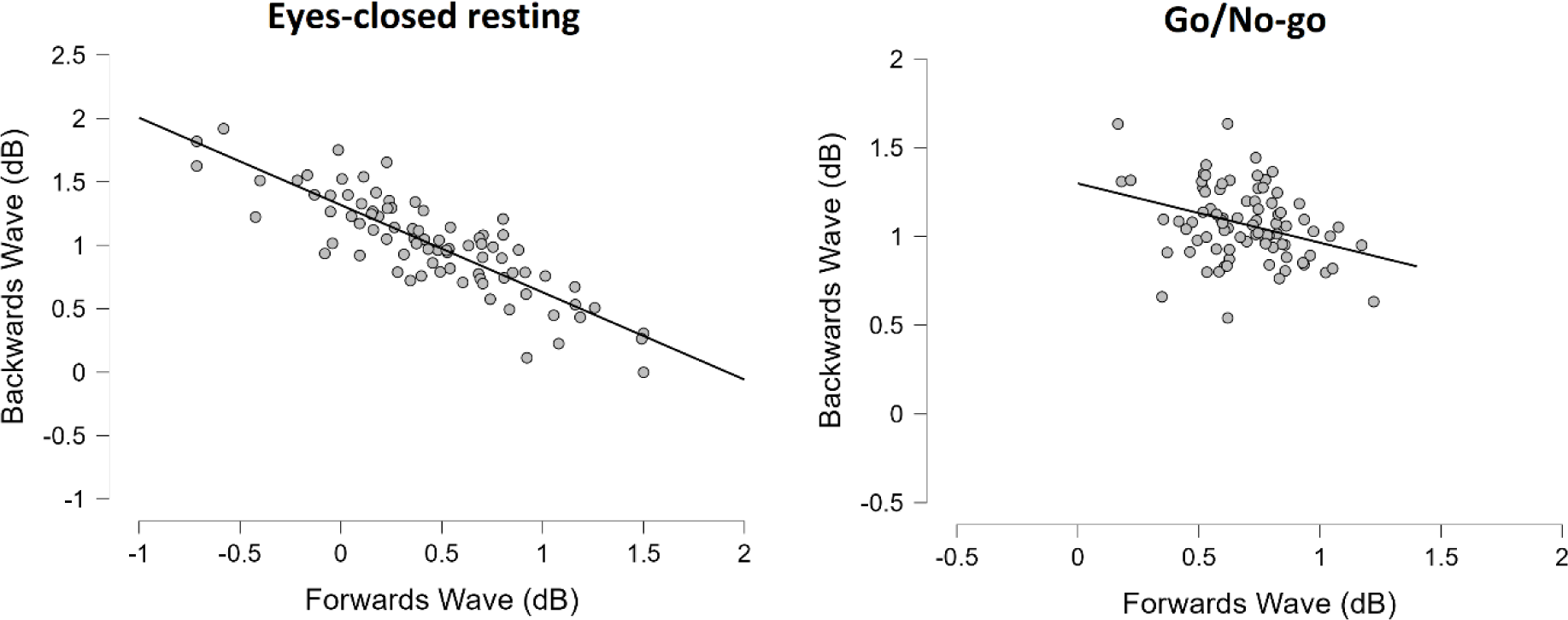
Correlations between forwards and backwards cortical travelling alpha wave strength in the eyes-closed resting EEG data and the Go/No-go task related EEG data. Note the broader spread of values (and different scale) in the eyes-closed resting data compared to the Go/No-go task related data, as well as the stronger correlation in the resting data (r = -0.848, 95% CI = - 0.774 to -0.899, p < 0.001, BF10 = 1.621*10^21^ and r = -0.333, 95% CI = -0.128 to -0.511, p = 0.001, BF10 = 15.079 respectively).

**Table 1.** Means and standard deviations (SD) for the strength of the forwards and backwards waves in the eyes-closed resting data for each group.

|  | Forwards Waves (dB) | Backwards Waves (dB) |
| --- | --- | --- |
|  | Mean (SD) | Mean (SD) |
| Meditators | 0.568 (0.487) | 0.922 (0.380) |
| Non-meditators | 0.358 (0.389) | 1.091 (0.291) |

### Meditators show stronger forwards travelling waves only during an attention task

While participants were undertaking a visual attention task, the strength of their forwards waves increased compared to while they were resting with their eyes closed. However, in alignment with the group differences in forwards travelling wave strength during resting, meditators again showed stronger forwards waves during the task. These results suggests that increased sensory processing is required to perform visual attention tasks compared to resting with the eyes closed, and that the higher trait forwards wave strength shown by meditators during resting allowed more effective increases in their forward wave strength during task demands. However, in contrast to the eyes-closed resting results, meditators showed a similar backwards wave strength during the attention task compared to non-meditators. This reflects an increase from meditator’s eyes-closed resting backwards wave strength to the attention task backwards wave strength. In contrast, non-meditators showed the same backwards wave strength while at rest and during the attention task. These results suggest that meditators were able to engage typical top-down attention processes when those processes were required for task completion, as well as continuing to generate stronger bottom-up sensory processing during the task. In contrast, non-meditators generated the same amount of top-down neural activity when resting with their eyes closed as they did while completing a cognitively demanding attention task.

With regards to Go/No-go task accuracy, the meditator group showed higher d-prime scores than the control group, indicating that meditators performed the task with higher accuracy across both Go and No-go trials types: t(124) = 2.751, p = 0.007, d = 0.492, BF10 = 5.554 (see Figure 5). Note that these results have been previously reported separately for sample 1 (Bailey, Freedman, et al., 2019), and are in preparation to be reported in a separate publication for sample 2.

**Figure 5.**
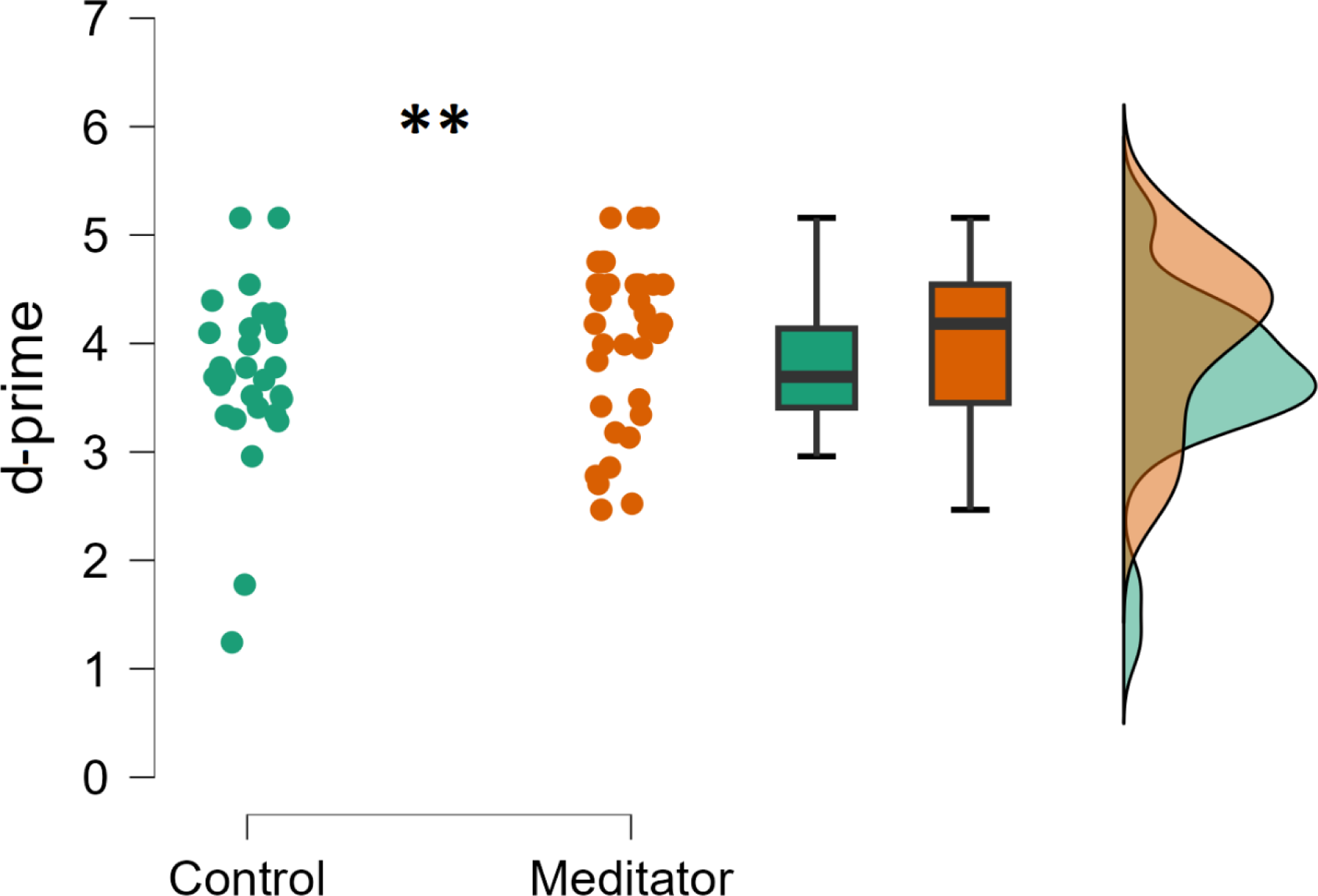
d-prime scores for each group in the Go/No-go task. The meditator group showed higher d-prime scores than the control group, suggesting higher task accuracy: t(124) = 2.751, p = 0.007, d = 0.492, BF10 = 5.554. ** p < 0.01.

In alignment with the resting data, the Go/No-go data showed a significant interaction between group and wave direction: F(1,124) = 6.059, p = 0.015, FDR-p = 0.015, ηp² = 0.047, ηG² = 0.033, BFincl = 11.068 (see Figure 6). Post-hoc t-tests showed that the interaction was driven by the meditators showing higher forwards wave strength than the non-meditators (FDR-p = 0.008, Cohen’s d = 0.518, BF10 = 7.942), but no significant differences in the backwards waves (FDR-p = 0.228, Cohen’s d = 0.217, BF10 = 0.371). There was also a significant main effect of wave direction, with both groups showing higher values for backwards wave strength than forwards wave strength: F(1,124) = 159.121, p < 0.001, ηp² = 0.562, ηG² = 0.470, BFincl = 2.217*10^32^. The main effect of group was not significant: F(1,124) = 2.173, p = 0.143, ηp² = 0.017, ηG² = 0.005, BFincl = 0.287.

**Figure 6.**
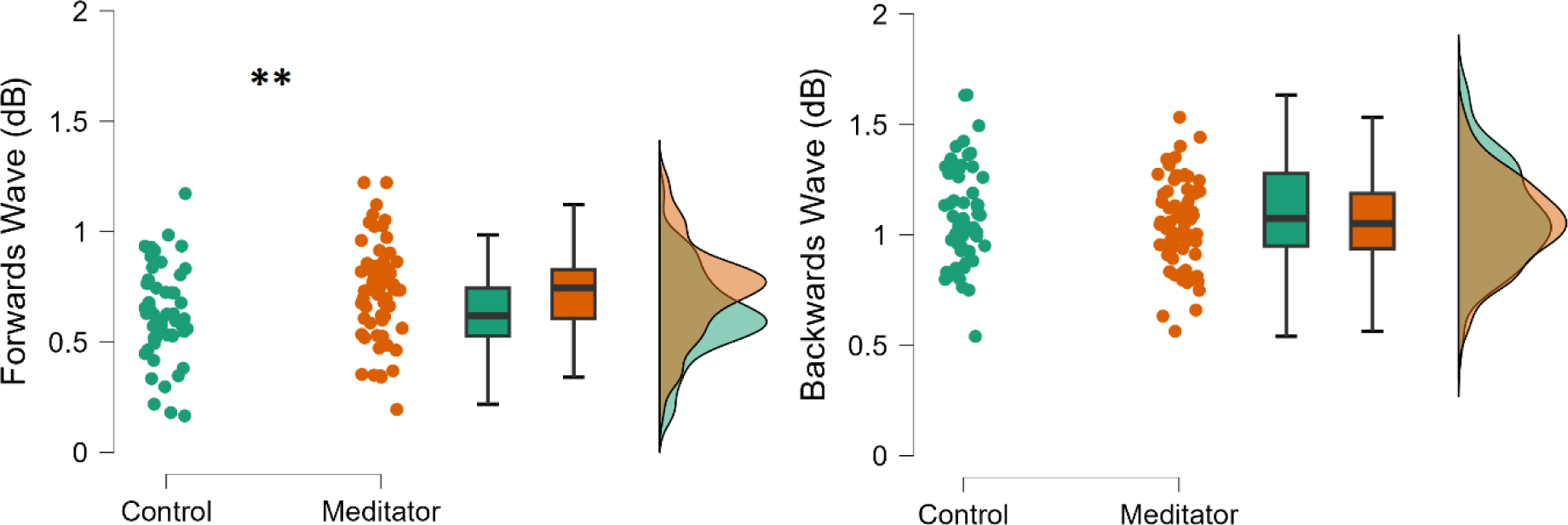
Forwards and backwards wave strength during the Go/No-go task within each group. Post-hoc t-tests showed that meditators had higher forwards wave strength than non- meditators (FDR-p = 0.008, Cohen’s d = 0.518, BF10 = 7.942), but no significant differences were present between groups in backwards wave strength (FDR-p = 0.228, Cohen’s d = 0.217, BF10 = 0.371). ** FDR-p < 0.01.

Additionally, while there was a negative correlation across participants between the strength of the forwards and backwards waves in the Go/No-go dataset, the confidence intervals for the r value suggested the correlation was significantly weaker than for the resting data (r = -0.333, 95% CI = -0.128 to -0.511, p = 0.001, BF10 = 15.079, compared to r = -0.848, 95% CI = -0.774 to - 0.899 for the eyes-closed resting data). Note that we restricted this correlation to only the participants who provided data for both the eyes-closed resting recordings and the Go/No-go task to enable direct comparison of the correlation strength between the two recording types. However, the entire Go/No-go dataset showed a very similar correlation strength (r = -0.393, 95% CI = -0.234 to -0.531, p = 0.001, BF10 = 3075.283). Finally, despite the difference between groups in both behavioural performance and in forwards wave strength in the Go/No-go task, no correlations were present across participants between the forwards or backwards travelling wave strength and task performance (all p > 0.2, all BF10 < 0.3).

**Table 2.** Means and standard deviations (SD) for the strength of the forwards and backwards waves in the Go/No-go data for each group.

|  | Forwards Waves (dB) | Backwards Waves (dB) |
| --- | --- | --- |
|  | Mean (SD) | Mean (SD) |
| Meditators | 0.732 (0.210) | 1.047 (0.197) |
| Non-meditators | 0.625 (0.201) | 1.093 (0.226) |

Finally, because the means from the resting and Go/No-go analyses suggested a potentially interesting interaction between group and resting/task for the backwards waves, we performed an exploratory repeated measures ANOVA restricted to just participants who had provided EEG data for both the eyes-closed resting and Go/No-go task. The analysis showed a significant interaction between group and recording condition: F(1,82) = 6.381, p = 0.013, ηp² = 0.072, ηG² = 0.026, BFincl = 3.795 (see Figure 7). Post-hoc tests indicated the interaction was driven by the meditators showing an increase in backwards wave strength from resting to the Go/No-go task (FDR-p = 0.030, Cohen’s d = 0.370, BF10 = 2.783), while controls showed no difference (FDR-p = 0.293, Cohen’s d = 0.176, BF10 = 0.299). In contrast, the interaction between group and task was not significant for the forwards waves: F(1, 82) = 1.256, p = 0.266, ηp² = 0.015, ηG² = 0.006, BFincl = 0.373. Instead, as suggested by the analysis of the Go/No-go and eyes-closed resting datasets independently, there was a significant main effect of group, with the meditators showing stronger forwards waves in both datasets: F(1, 82) = 7.388, p = 0.008, ηp² = 0.083, ηG² = 0.052, BFincl = 4.537.

**Figure 7.**
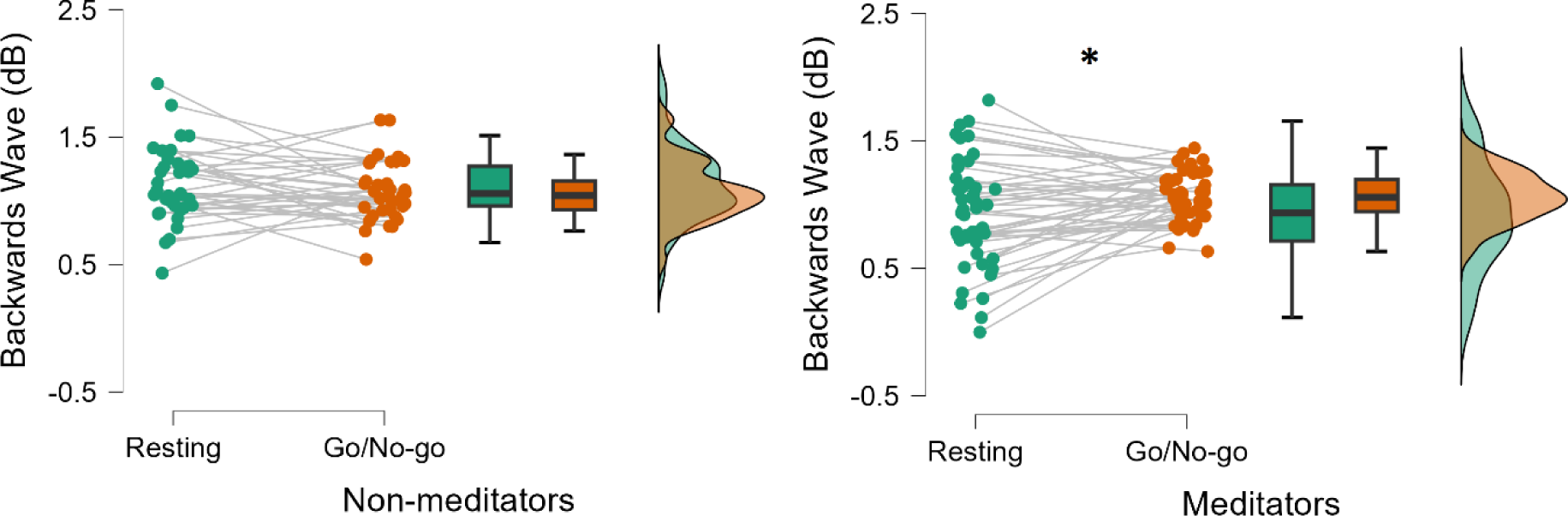
Backwards waves within each group from the eyes-closed resting and Go/No-go datasets. There was a significant interaction between group and resting/task: F(1,82) = 6.381, p = 0.013, ηp² = 0.072, ηG² = 0.026, BFincl = 3.795. Post-hoc t-tests indicated the interaction was driven by the meditators showing an increase in their backwards wave strength from resting to the Go/No-go task (FDR-p = 0.030, Cohen’s d = 0.370, BF10 = 2.783), while controls showed no difference (FDR-p = 0.293, Cohen’s d = 0.176, BF10 = 0.299). * FDR-p < 0.05.

### Travelling wave strengths are modulated by working memory processes

Our analysis of travelling waves during the working memory task showed that forwards travelling wave strength was higher during memory set presentation, during which time participants processed visual information, attempting to remember it for later recall. In contrast, backwards wave strength was higher while the probe stimuli were being presented, during which time participants had to respond as to whether the probe stimulus matched their recollection of the memory set stimuli. These results provide support for suggestions that forwards travelling alpha waves are indicative of an underlying function that encodes visual information (Lozano-Soldevilla & VanRullen, 2019; Mohan et al., 2024), and that backwards travelling waves may act as top-down modulation of precision to reinstate the cortical representations of memory set items required for successful memory retrieval (Barron et al., 2020; Mohan et al., 2024).

Within the Sternberg working memory task data, meditators showed higher accuracy, a result that we have reported previously (Bailey et al., 2020): t(51) = 2.503, p = 0.008, Cohen’s d = 0.688, BF10 = 3.373. Additionally, our travelling wave analysis showed there was a significant interaction between the period of the task and the direction of the wave: F(2,112) = 5.041, p = 0.008, ηp² = 0.083, ηG² = 0.030, BFincl = 140.688 (see Figure 8). Post-hoc t-tests indicated the forward wave strength was significantly greater in the memory set period compared to the delay period (FDR-p = 0.006, Cohen’s d = 0.502, BF10 = 74.415) and compared to the probe period (FDR-p = 0.015, Cohen’s d = 0.350, BF10 = 3.533), while the delay and probe periods did not differ from each other (FDR-p = 0.059, Cohen’s d = 0.264, BF10 = 0.939). In contrast, backwards wave strength was significantly higher in the probe period compared to the memory set period (FDR-p = 0.006, Cohen’s d = 0.422, BF10 = 13.605), and compared to the delay period (FDR-p = 0.015, Cohen’s d = 0.356, BF10 = 3.953), while the memory set period and the delay period did not differ from each other (FDR-p = 0.903, Cohen’s d = 0.016, BF10 = 0.145). We note that in addition to differences in working memory processes required by the different task periods, the probe periods contained a motor response while the other periods did not. To test whether the motor response might explain the differences between the probe period and other working memory periods, we performed an exploratory analysis of the Go/No-go task, which contained trials both with and without motor responses. We compared travelling wave strength within the Go and No-go trials separately, as Go trials contained a motor response (similar to the probe period of the working memory task) and the No-go trials did not (similar to the memory set and delay periods of the working memory task). This analysis (reported in full in the Supplementary Materials) showed no difference between the Go and No-go trials, suggesting the results related to the probe period of the working memory task were unlikely to be simply due to the presence of a motor response.

**Figure 8.**
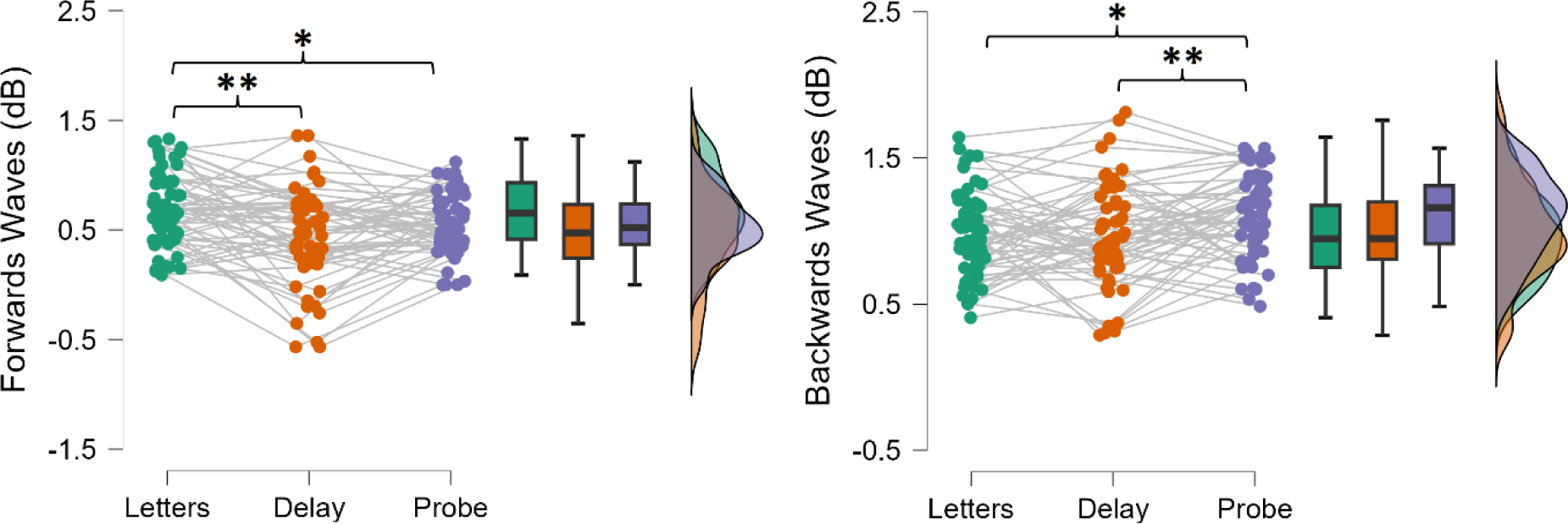
Forwards and backwards wave strength (in decibels – dB) for each period of the Sternberg working memory task across both groups. Post-hoc t-tests indicated forward wave strength was significantly greater in the memory set period (“Letters”) compared to the delay period (FDR-p = 0.006, Cohen’s d = 0.502, BF10 = 74.415). Forwards wave strength was also greater in the memory set period compared to the probe period (FDR-p = 0.015, Cohen’s d = 0.350, BF10 = 3.533), while the delay and probe periods did not differ from each other (FDR-p = 0.059, Cohen’s d = 0.264, BF10 = 0.939). In contrast, backwards wave strength was significantly higher in the probe period compared to the memory set period (FDR-p = 0.006, Cohen’s d = 0.422, BF10 = 13.605), and compared to the delay period (FDR-p = 0.015, Cohen’s d = 0.356, BF10 = 3.953), while the memory set period and the delay period did not differ (FDR-p = 0.903, Cohen’s d = 0.016, BF10 = 0.145). ** FDR-p < 0.01. * FDR-p < 0.05.

In contrast to our results for the resting and Go/No-go data, our analysis of the working memory task showed no significant main effects or interactions involving group (all p > 0.09, all BF10 < 0.4, reported in full in the supplementary materials). It is possible that this null result was due to the smaller sample size for these comparisons, with the Bayes Factor providing inconsequential evidence for the null hypothesis for some of the interactions involving group, and the between group patterns showing the same directions as the eyes-closed resting and Go/No-go task results (see the Supplementary Materials and Supplementary Materials Figure 1 for an exploration of this point).

Results of the regression to test associations between wave strength and working memory performance indicated that the overall regression model was significant: F(7, 50) = 2.262, p = 0.044, with a model that included forwards wave strength in both the memory set presentation period and probe period providing evidence in support of the alternative hypothesis: BF10 = 8.468. The effects were driven by a significant interaction, where forward wave strength in the memory set presentation period was related to better d-prime scores (t = 2.234, p = 0.030), while forward wave strength in the probe presentation period was related to worse d-prime scores (t = -2.595, p = 0.012). Group was also a significant predictor (t = 2.105, p = 0.040). This effect of group aligns with the higher working memory accuracy from meditators shown in our previous research on this dataset (Bailey et al., 2020). See Figure 9 for a depiction of the marginal effects of forwards waves in the memory set presentation and probe periods on d- prime scores.

**Figure 9.**
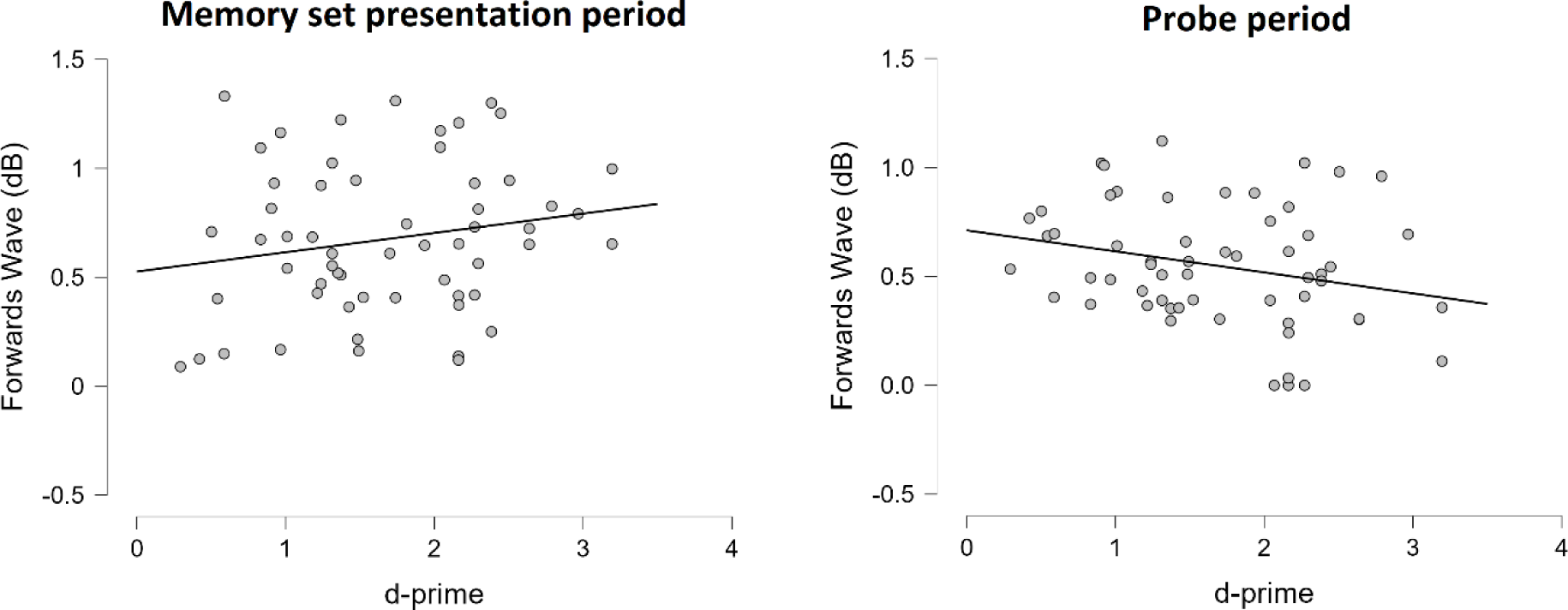
Regression analysis indicated that, controlling for other factors, performance in the working memory task was predicted by forward wave strength in both the memory set presentation period and the probe presentation period. Results showed forward wave strength in the memory set presentation period predicted better d-prime scores (t = 2.234, p = 0.030), while forward wave strength in the probe presentation period predicted worse d-prime scores (t = -2.595, p = 0.012).

**Table 3.** Means and standard deviations (SD) for the strength of the forwards and backwards travelling cortical alpha waves in the Go/No-go data for each group.

|  |  | **Meditators** | **Non-meditators** |
| --- | --- | --- | --- |
|  |  | **Mean (SD)** | **Mean (SD)** |
| **Forwards Waves (dB)** | Memory set presentation period | 0.739 (0.331) | 0.610 (0.356) |
|  | Delay period | 0.478 (0.418) | 0.401 (0.463) |
|  | Probe presentation period | 0.567 (0.293) | 0.530 (0.261) |
| **Backwards Waves (db)** | Memory set presentation period | 0.951 (0.310) | 1.000 (0.285) |
|  | Delay period | 0.999 (0.351) | 0.964 (0.348) |
|  | Probe presentation period | 1.151 (0.306) | 1.062 (0.265) |

## Discussion

Our results showed that experienced mindfulness meditators generate stronger forwards travelling cortical alpha waves compared to non-meditators, both while resting with their eyes-closed and while performing a demanding visual attention task. Furthermore, meditators showed weaker backwards travelling cortical alpha waves while resting with their eyes-closed compared to non-meditators. Meditators also showed an increase in the strength of their backwards waves from eyes-closed resting to the visual attention task, while non-meditators showed no difference in backwards wave strength between eyes-closed resting and the task. Finally, the forwards and backwards travelling waves were differently modulated during different periods of a working memory task and some of these modulations were associated with working memory performance. These results have important implications for our understanding of the neural effects of meditation, providing indications of specific neural mechanisms underlying the effects of meditation, and providing support for a predictive processing explanation of how meditation alters brain activity to produce changes in subjective experience and cognitive function. The results also have broader implications for our general understanding of brain function.

### Implications for our understanding of the effects of meditation

Previous research has indicated that meditators show a greater ability to modulate standing alpha oscillatory power to meet task demands, with meditators showing a larger capacity to increase alpha power in visual regions when a task presents only somatosensory stimuli, and a larger capacity to increase alpha power in somatosensory regions when visual stimuli are presented (Wang et al., 2020). We have also shown that meditators are able to decrease alpha power within specific cognitive task-relevant time periods and to increase alpha phase synchronisation to stimuli presentation timing (Bailey et al., 2023a). The current results extend these previous results regarding standing wave modulation to travelling waves.

Forwards travelling cortical alpha waves are generated in response to the presentation of a stimulus, and are thought to be indicative of an underlying mechanism for processing sensory information (Pang et al., 2020). As such, the higher strength of forwards waves in meditators aligns with an important component of meditation practice, which trains the continual direction of attention to sensations (including redirection of attention back to sensations when attention lapses). Previous research has also suggested that forwards waves reflect sensory predictive processing errors being passed up to progressively higher layers in the cortical hierarchy (Alamia & VanRullen, 2019). As such, our results align with a predictive processing account of the effects of meditation, which suggests that the repeated allocation of attention to sensations produces neuroplastic changes that increase the synaptic gain (and therefore precision) with which bottom-up prediction errors are processed (Laukkonen & Slagter, 2021; Lutz et al., 2019; Manjaly & Iglesias, 2020). In contrast, backwards travelling waves are spontaneously generated, and have been suggested to index the strength of higher-order predictions generated by frontal brain regions (Alamia & VanRullen, 2019; Pang et al., 2020). As such, the lower strength of backwards waves during eyes-closed resting in the meditator group also aligns with an important component of meditation practice, with its focus on training nonjudgemental present moment awareness of current sensations (Kabat-Zinn, 2023). The lower strength of backwards travelling cortical wave strength in meditators also provides support for a predictive processing account of meditation, which suggests that the practice of non-judgement present-moment awareness produces a trait reduction in the generation and elaboration of higher-order predictions (Deane et al., 2020; Lutz et al., 2019). Our findings regarding backwards wave strength also suggest that the effects of long-term meditation are not driven by an increase in top-down attentional control processes (Garland et al., 2009), which would be reflected by stronger backwards travelling waves. Instead, our results suggest the effects of meditation are likely more driven by increases to bottom-up processing (Chambers et al., 2009; Chiesa et al., 2013).

Interestingly, meditators showed an increase in the strength of their backwards travelling waves from eyes-closed resting (where they showed lower backwards travelling wave strength than non-meditators) to performance of the visual attention task (where they showed the same backwards travelling wave strength as non-meditators). In contrast, non-meditators showed no difference between the eyes-closed resting and task conditions. It is worth noting that in addition to the visual processing requirements of the Go/No-go task, higher-order cognitive functions were critical for accurate performance of the task. These higher-order functions include response inhibition, performance monitoring, and working memory to remember task demands. Additionally, since no more than three Go or No-go trials were presented consecutively, the sequence of stimuli presentation was weakly predictable, so sequence pattern recognition and prediction of upcoming stimuli may have also been useful for accurate task performance. As such, accurate task performance likely required the generation of higher-order predictions, which are suggested to be indexed by backwards travelling waves, in addition to visual processing indexed by the forwards waves (Alamia & VanRullen, 2019; Pang et al., 2020).

In this context, it is perhaps more interesting to explore the finding that non-meditators generated the same level of backwards waves during eyes-closed resting as they did during the cognitive task. This result suggests that the non-meditators engaged in similar levels of higher-order cognitive activity while their brain was simply resting, with no specific task demands, as they did while performing demanding executive functions. Our suggestion is that the high level of backwards waves during eyes-closed resting in non-meditators reflects the continual generation and elaboration of higher-order predictions, even in the absence of task requirements. These higher-order predictions likely consist of thoughts or imagery about oneself, the future or the past (Smallwood & Schooler, 2006; Stawarczyk et al., 2011), which may be brought about through the instantiation of counterfactual or ‘fictive’ prediction errors in the absence of task demands (Barron et al., 2020; Immink & Corcoran, 2023; Lutz et al., 2019). Viewed from this perspective, the lower strength of backwards waves shown by meditators during eyes-closed resting may be a marker of the reductions in rumination and worry that are suggested as a psychological mechanism by which mindfulness improves well-being and protects against depressive relapse (van der Velden et al., 2015).

In contrast, the finding of stronger forwards travelling waves from occipital (visual processing) brain regions in meditators during eyes-closed resting (i.e., in the absence of visual stimulation), is not as intuitive to explain. Previous research has indicated that while top-down attention modulates backwards waves irrespective of visual stimuli, forwards waves are only modified by attention in the presence of visual stimulation (Alamia, Terral, et al., 2023). This poses an obvious question - why would meditators generate stronger forwards waves (which are associated with visual processing) when no visual stimuli are presented? To offer one potential explanation, we note that a common instruction from the Vipassana meditation tradition is to focus on perceiving reality “as it is”, not as it is desired to be (Goenka, 2012). By following this instruction, it may be that meditators are processing the absence of external visual stimulation “as it is” in a similar manner to the way they would process the presence of external visual stimulation. That is, meditators may simply be processing the experience that “it is dark when I have my eyes closed” with increased synaptic gain compared to non-meditators.

Alternatively, the result could be explained by the fact that the majority of the meditators had spent years training their attention to sensations, enhancing the precision-weighting and synaptic gain of sensory brain regions (Laukkonen & Slagter, 2021; Lutz et al., 2019; Manjaly & Iglesias, 2020). This effect may be maintained even in the absence of visual stimulation. Additionally, while the travelling waves were measured from occipital electrodes to frontal electrodes, it may be that auditory and interoceptive processing also modulated the travelling waves, with the sounds of the participant’s breath and bodily sensations present even when participants rested with their eyes closed. Perhaps meditators were devoting more neural processing to these sensations, while non-meditators devoted more neural resources to processing autobiographical or future-planning mental content. A third explanation might be that the greater strength of forwards waves is simply the result of the lower strength of backwards waves in the meditators, due to the tight coupling between the two wave directions (Pang et al., 2020) and the negative feedback loop between brain regions generating predictions and brain regions passing prediction errors up the neural hierarchy (Friston, 2019). However, comparing the correlations between forwards and backwards waves separately in the resting and task related data suggests the inverse relationship between forwards and backwards waves is not causal or immutable. As such, this explanation might be the least likely of these three explanations to be accurate.

Irrespective of the explanation for the higher strength of resting forwards waves in meditators, we note that meditators showed a similar increase to the non-meditators in their forwards wave strength from eyes-closed resting to the cognitive task. Since the meditators showed a higher eyes-closed resting forwards wave strength, which might be considered to reflect a “baseline” state, it might be that the higher baseline forwards waves strength shown by meditators allowed for even stronger generation of forwards waves during visual attention tasks, perhaps indicating an increased capacity for visual processing that aligns with better task performance. However, it is also worth noting that the difference in wave strength between meditators and non-meditators might not be ubiquitous across all situations, as no difference was detected between the groups in the working memory task. This is also in alignment with previous research, which has suggested that differences in neural activity between meditators and non-meditators are present in some tasks, but not others (Bailey et al., 2023a). Having said that, we note that the pattern of stronger forwards wave in meditators was present (but not significant) in the working memory task, and that our Bayesian analysis did not provide evidence to support the null hypothesis. As such, a larger sample size may be required to detect differences between meditators and non-meditators in travelling waves related to working memory tasks.

Alternatively, we speculate that one potential explanation for the presence of a difference in the Go/No-go task but absence in the working memory task could be that the rapid stimuli presentation of the Go/No-go task prompting stronger engagement of attentional mechanisms, and this extra task demand revealed the increased ability in meditators to modulate forwards waves to meet task demands. In contrast, the slower display of the working memory stimuli may have not required the same engagement of attentional mechanisms, and so the lower task demand meant that the increased ability of meditators to modulate forwards waves was not revealed by the working memory task. It may be interesting for future research to explore which contexts the higher forwards wave strength is present in meditators.

It is also interesting to speculate about the changes to neuroanatomy that might enable the alterations to travelling waves that we have observed in meditators, and whether travelling waves might be associated with specific functions that are altered by meditation. While our research demonstrates differences in forwards and backwards travelling wave strength in meditators, and previous research has suggested these travelling waves likely relate to predictive processing mechanisms, the mechanisms underlying our results are not obvious. Alpha oscillations have been demonstrated to be propagated by intra-cortical axonal connections (Hindriks et al., 2014). To produce travelling waves, these intra-cortical axonal connections need to connect neuron populations that are intrinsically oscillatory, with the connections between these populations being weak so that each population influences the other population but cannot distort the oscillatory cycle length produced by other neuronal populations (Ermentrout & Kleinfeld, 2001). As such, one potential candidate mechanism to explain our results might be a progressive neuroplastic strengthening of forwards projecting intra-cortical axonal connections with repeated meditation practice, which over time resulted in a baseline increase in the strength of forwards travelling cortical alpha waves.

Alternatively, research has suggested that alpha synchronisation across cortical regions is coordinated by deeper cortical layers (Alamia & VanRullen, 2024). This coordination by the deeper layers might enable the synchronisation of more superficial layers, which then generate higher oscillatory frequencies to facilitate information processing (Alamia & VanRullen, 2024). In particular, it has been suggested that thalamic nuclei, including the pulvinar and reticular nuclei are generators of alpha oscillations, and are broadly connected to cortical regions at multiple hierarchical levels (Fiebelkorn & Kastner, 2019; Saalmann et al., 2012). The pulvinar is also implicated in attention and working memory function, so has been suggested to be a potential candidate region that enables coordination of travelling waves (Alamia & VanRullen, 2024). It may be that meditation training increases the ability of the pulvinar to regulate cortical travelling waves, through strengthening of feedforward and feedback loops between the thalamus and cortex. In this context, we note there is preliminary evidence that thalamocortical functional connectivity is enhanced by meditation, both from studies using functional magnetic resonance imaging and studies using EEG data combined with modelling (Chen et al., 2021; Saggar et al., 2015). Furthermore, the study using EEG and modelling showed that the enhanced thalamocortical functional connectivity was associated with increased dynamic stability of the EEG activity (Saggar et al., 2015). Applying a comprehensive data-driven approach to analyse the resting EEG data reported in the current study, we also found that meditators had higher dynamic stability of their EEG activity compared to the non-meditators (Bailey et al., 2024).

Regardless of the specific mechanisms that drive alterations in travelling waves, the function of travelling waves might be to enable sensitisation of neurons to produce action potentials, due to the associated voltage shifts in the extracellular material which make neurons more likely to depolarize (von Wegner et al., 2021). If the brain generates travelling waves so that predictions or prediction errors processed within a specific cortical region arise in that region with timing aligned to an oscillatory phase that provides sensitisation, then there may be an increase in the likelihood for a neuron to depolarize, and thus an increase in the processing of the prediction or prediction error (Alamia & VanRullen, 2024). As such, an increased ability to modulate travelling cortical alpha waves may reflect an attentional mechanism enabling increased perceptual sensitivity and more finely tuned top-down attentional control. However, further research is required to determine the exact physiological origins and functions of the travelling wave effect measured in scalp EEG data.

### Implications for our understanding of the brain, predictive processing, and potential clinical applications

To allow adaptive behaviour, the brain needs to be able to both coordinate neural activity across brain regions and networks and adjust the coordination to meet the demands of the current environment (Mohan et al., 2024). Alpha oscillations seem to provide some of these coordination functions. Previous research on the function of standing alpha oscillations has indicated that alpha activity is primarily associated with two attention-related functions. The first of these is an active top-down inhibitory function that allows frontal brain regions to inhibit sensory processing regions when those regions are not relevant to current goals (Klimesch, 2012). The second function is a timing function, where specific alpha oscillatory phases can be timed to match the processing of incoming stimulus information, increasing the likelihood of neuronal activation in response to those stimuli (Klimesch, 2012). Analogues of these two functions have also been demonstrated when examining the function of travelling alpha waves, where top-down attention has been shown to modulate backwards waves irrespective of whether visual stimuli are presented, while forwards waves have been demonstrated to only be modified by attention in the presence of visual stimulation (Alamia, Terral, et al., 2023). Our results provide further evidence for travelling waves as a correlate of large-scale network coordination. In particular, our results provide additional support for an association between forwards waves and visual attention (Alamia, Terral, et al., 2023; Lozano-Soldevilla & VanRullen, 2019; Pang et al., 2020), with both groups showing an increase in forwards wave strength from eyes-closed resting to the visual cognitive task. However, in contrast to previous research indicating that visual stimulation decreases backwards waves (Alamia et al., 2020), our results also showed that backwards waves could be maintained at the same level (in non-meditators), or even increased (in meditators) in a visual attention task that also required higher-order processes for accurate task performance. This provides support for a previously untested suggestion from the literature that backwards waves are important when top-down expectations are involved in a task (Alamia & VanRullen, 2024).

Our results also show that the modulation of forwards and backwards travelling wave strength is associated with specific periods of a working memory task, where specific working memory processes are required for accurate task performance. In particular, the memory set presentation period where participants attempted to encode stimuli for later recall was associated with higher forward wave strength. Higher forwards wave strength during this period predicted task accuracy across participants, providing further support for research suggesting that forwards travelling alpha waves index an underlying mechanism that functions to encode visual information (Lozano-Soldevilla & VanRullen, 2019; Mohan et al., 2024). In contrast, the working memory probe presentation period (where participants attempted to recall whether the probe stimulus was in the memory set that they were presented earlier) was associated with an increase in backwards waves. This increase in backwards waves is perhaps indicative of a top-down modulation of precision to reinstate cortical representations of memory set items enabling determination of whether they matched the probe stimulus (Barron et al., 2020; Mohan et al., 2024). Across participants, our results also provided further support for links between forwards waves and successful encoding (Mohan et al., 2024). However, interestingly, our results indicated that lower forward wave strength during the probe period of the working memory task was related to better task performance. We measured wave strength after the initial period of visual processing, so stronger forwards waves at that time may reflect distraction by visual stimuli when top-down working memory processes were required for accurate task performance, perhaps explaining this negative relationship.

Our results also provide the first evidence that forwards and backwards wave strengths are likely to be modifiable by an attention training technique. In particular, our results indicate that attention training to sensations is associated with increases in the strength of forwards travelling waves, which have been interpreted as indexing a mechanism for the processing of bottom-up sensory information (Pang et al., 2023), reflecting the feedforward passing of hierarchical prediction errors (Alamia & VanRullen, 2019). Additionally, meditators showed lower strengths of backwards waves during eyes-closed resting conditions, but not during a cognitive task. This result suggests that the typical generation or elaboration of higher-order predictions that typically occurs while at rest might be reducible, potentially by the present moment focus and non-judgemental aspects of meditation practice, in a manner that does not also reduce the potential to engage higher-order prediction generation when environmental demands require. This result might be particularly valuable for extension to clinical applications. In particular, recent research has shown that individuals with schizophrenia display elevated levels of backwards travelling waves (Alamia, Gordillo, et al., 2023), and individuals with post-traumatic stress disorder (PTSD) show lower posterior to frontal alpha functional connectivity (which might align with reduced forwards travelling waves) (Clancy et al., 2018). The same research indicated that when this impaired functional connectivity was increased by neuromodulation, PTSD symptoms were improved, suggesting a potential for meditation practice to have the same effect (Clancy et al., 2018).

Previous research has also shown that forwards and backwards wave strength are negatively correlated, so that within each participant, when backwards wave strength is increased, forwards wave strength decreases (Alamia et al., 2020). Our results provide support for this finding, showing that forwards and backwards travelling wave strength were strongly negatively correlated across individuals. However, our results also indicate that across individuals, this relationship was much stronger in the eyes-closed resting data than in the task data.

Furthermore, while visual stimulation without a task has been shown to prompt an increase in forwards waves and concurrent decrease in backwards waves that preserves the strong inverse relationship between the two wave directions (Pang et al., 2020), when shifting from the eyes-closed resting to the visual attention task, meditators showed an increase in forwards wave strength *and* an increase in backwards wave strength. This suggests that the relationship between the wave directions is not immutable, and that in our data, the shift from eyes-closed resting to task demands may have elicited an increase in both forwards waves (to process the visual information) and an increase in backwards wave strength (in meditators) to enable task-relevant predictive processing.

Interestingly, a similar pattern of results to the increased forwards and decreased backwards wave strength in meditators has been shown in a study of the effects of the psychedelic drug Dimethyltryptamine (DMT) (Alamia et al., 2020). However, despite the superficial similarity, we suggest that the effects of DMT are clearly driven by different mechanisms to those associated with meditation. In particular, DMT is a mixed serotonin receptor agonist that induces prominent visual hallucinations (Alamia et al., 2020). As such, the increase in forwards waves and decrease in backwards waves associated with DMT likely reflects “visual processing” in the absence of actual visual stimuli, driven by a pharmacological perturbation of the typical function of the visual cortex (Alamia et al., 2020). In contrast, even after extensive meditation experience, visual hallucinations are rare (Schlosser et al., 2019). As such, the higher strength of forwards waves in meditators is unlikely to reflect the processing of visual hallucinations. Instead, the meditators in our study had spent years (on average) training their attention to sensations, enhancing the precision-weighting and synaptic gain of sensory brain regions, even in the absence of sensory stimulation, reflecting an increase in sensory processing even in the absence of visual stimuli. Additionally, we note that the effects of DMT also induce a marked reduction in alpha power, whereas our research shows that alpha power increases are associated with meditation (McQueen et al., 2023). As such, our results should not be taken as an indication that the DMT experience is similar to the experience of having significant experience with meditation practice.

### Limitations and future directions

While our results provide several interesting findings, our study is not without limitations. First, our analyses were limited to only examining travelling waves from posterior to frontal electrodes and vice versa along the midline. Rotating waves are also present in the cortex (Ermentrout & Kleinfeld, 2001), and these rotating waves are likely to also be linked to important brain functions, with recent research suggesting they might provide a more complete explanation of the microstate patterns that are consistently found in human EEG activity (von Wegner et al., 2021). We also note that while we have referred to travelling waves throughout this manuscript, the phenomena that appears as travelling waves in MEG data may be explicable instead by activation of an occipital source dipole followed by a phase-lagged activation of a parietal source dipole (Zhigalov & Jensen, 2023). Additionally, traveling waves detected in scalp EEG electrodes show different properties to those measured directly from the cortical surface, and it has been suggested that scalp waves might be generated by progressive activation of individual source generators of brain activity (Orczyk & Kajikawa, 2022). As such, our results might reflect the sequential activations of dipoles, rather than a cortical travelling wave per se. However, the activation of occipital and parietal dipoles are unlikely to explain backwards travelling waves which originate in frontal regions (Alamia, Terral, et al., 2023). We also note that our working memory results show a strong alignment with the results of analyses performed using intracortical electrodes where travelling waves can be more strongly verified (Mohan et al., 2024). Finally, we note that given the breadth of electrode coverage included in our analysis (midline electrodes from occipital to fronto-polar regions), our results likely pertain to macroscopic travelling waves, with waves that propagate between brain regions, and do not speak to travelling waves at smaller scales (Alamia & VanRullen, 2024). As such, more research is required to examine travelling waves in other directions and locations, and the underlying mechanisms of the travelling waves.

Second, there were no correlations between Go/No-go task performance and travelling wave strength, querying the functional relevance of the travelling waves. However, we note that participants generally performed highly accurately in the Go/No-go task, so this may reflect a ceiling effect. Additionally, our analysis focused only on the strength of the travelling waves. Some previous research has suggested that a faster speed of forwards alpha travelling waves relates to reaction times in a visual image identification task (Fellinger et al., 2012), so potential differences between meditators and non-meditators in the speed of the cortical travelling waves might be worth exploring in future research. We also note that our analysis of travelling waves used the entire one second period after presentation of each Go and No-go stimulus, and our correlations tested travelling wave strength averaged across all trials within each participant. Forwards and backwards wave strengths have been shown to dynamically shift in response to task demands (Alamia, Terral, et al., 2023). As such, it may be that different time periods within each trial would have required engagement of different patterns of travelling waves. In particular, the initial presentation of stimuli might have elicited higher forward wave strength to process visual information, while the later period may have required an increase in backwards travelling wave strength to engage the executive functions enabling accurate responses. As such, a more fine-grained single trial analysis that separated early and late periods within each epoch may have revealed associations with behavioural performance. This proposal is supported by previous research which showed an association between increased forwards waves, visual attention, and task performance, suggesting it may be valuable for future research to explore different time periods for analysis (Alamia, Terral, et al., 2023).

On a related point, we note that the probe period of the Sternberg working memory task required a button response, while the other periods did not, posing a potential confound to our conclusion that modulation of travelling wave strength reflects the engagement of different working memory processes across the task. However, our exploratory analysis of the Go/No-go task (reported in full in the Supplementary Materials) indicated that travelling waves were not differently modulated by the Go and No-go trials (where one trial required a response, and the other did not). This suggests that the travelling wave modulation we detected in the working memory task was not simply driven by the presence of a button press response in the probe period and the lack of a motor response in the other periods, and instead was likely driven by differential engagement of working memory processes in alignment with previous research (Mohan et al., 2024).

Another obvious limitation is the use of a cross-sectional design, which does not allow for the inference of causality. However, a longitudinal study including participants with the same average degree of experience with meditation practice as the current study (an average of eight years since participants started practicing meditation) would be incredibly difficult to implement. Our results indicated only small to moderate effect sizes, detection of which was only made possible with our relatively large sample size. Longitudinal studies which typically include much less experience with meditation than the current study might be expected to produce even smaller effect sizes, making it less likely that longitudinal studies would detect significant differences between a meditation and control condition. Having noted this point, it may be interesting for future research to explore whether a reduction in eyes-closed resting backwards waves could be a marker of reduced rumination or worry following a mindfulness intervention for depression, as this has been suggested to be a mechanism of action of mindfulness interventions for depression (van der Velden et al., 2015). If this were the case, backwards travelling cortical alpha waves could be a useful indicator of the success of an intervention and might be usefully explored as a marker of the mechanism of action of meditation. If future research verifies our results, then travelling waves might also have utility as a method of providing feedback about an individual’s progress with meditation training, or even a potential predictor of which individuals might benefit most from a mindfulness intervention.

Finally, we explicitly provided participants with the instruction to only rest with their eyes closed and simply let their mind do as it would, but not to meditate. However, it is impossible to confirm that the meditation group did not simply use this time as an opportunity to meditate, or that they did not habitually commence their meditation practice during the eyes-closed resting recording. If the meditation group was practicing meditation instead of simply resting, our results might reflect neural activity during the state of meditation rather than trait differences in brain activity because of their meditation experience, in which case the finding that meditators showed weaker backwards waves might reflect the state of meditation rather than a trait difference between the groups. However, meditators also showed stronger forwards waves during the Go/No-go task, when they would not have been able to meditate, indicating that the greater strength of forwards waves (and associated increase in synaptic gain and precision of sensory processing) is a trait associated with long-term meditation experience. Additionally, in a sense, if the meditation group did habitually commence meditating instead of resting, this may be indicative of the natural habits of the meditator’s brain (given the instructions provided to participants). As such, the results still reflect the differences in brain activity and altered predictive processing-related neural activity associated with experience in meditation, regardless of whether they were resting with their eyes open or meditating.

To conclude, our results showed that experienced meditators generated stronger forwards travelling cortical alpha waves than non-meditators, both while resting with their eyes closed and during task performance. Meditators also generated weaker backwards travelling cortical alpha waves while resting but increased the strength of these backwards travelling waves to be equivalent to non-meditators during a visual attention task. These results align with a predictive processing perspective of the effects of meditation, with stronger forwards waves reflecting greater synaptic gain on ascending sensory prediction errors associated with attention training towards sensations, while weaker backwards waves might be indicative of less generation and elaboration of higher-order beliefs (or less mind-wandering) when no task demands are present.

## Ethics Information

Ethics approval for the first dataset was provided by the Ethics Committees of Monash University and Alfred Hospital.

## Data Availability Statement

The data that support the findings of this study are available from the corresponding author, NWB, upon reasonable request.

