## Supplementary Materials for "Experienced meditators show greater forward travelling cortical alpha wave strengths"

Within the Sternberg working memory task, there was no significant main effect of group: $F(1, 56) = 2.898$, $p = 0.094$, $\eta p^2 = 0.049$, $\eta G^2 = 0.006$, BFincl = 0.397. Nor was there a significant interaction between group and task period: $F(1, 56) = 0.155$, $p = 0.857$, $\eta p^2 = 0.003$, $\eta G^2 < 0.001$, BFincl = 0.078, nor between group and direction: $F(1, 56) = 0.223$, $p = 0.639$, $\eta p^2 = 0.004$, $\eta G^2 = 0.002$, BFincl = 0.348, nor between group, task period and direction: $F(1, 56) = 0.807$, $p = 0.449$, $\eta p^2 = 0.014$, $\eta G^2 = 0.005$, BFincl = 0.367.

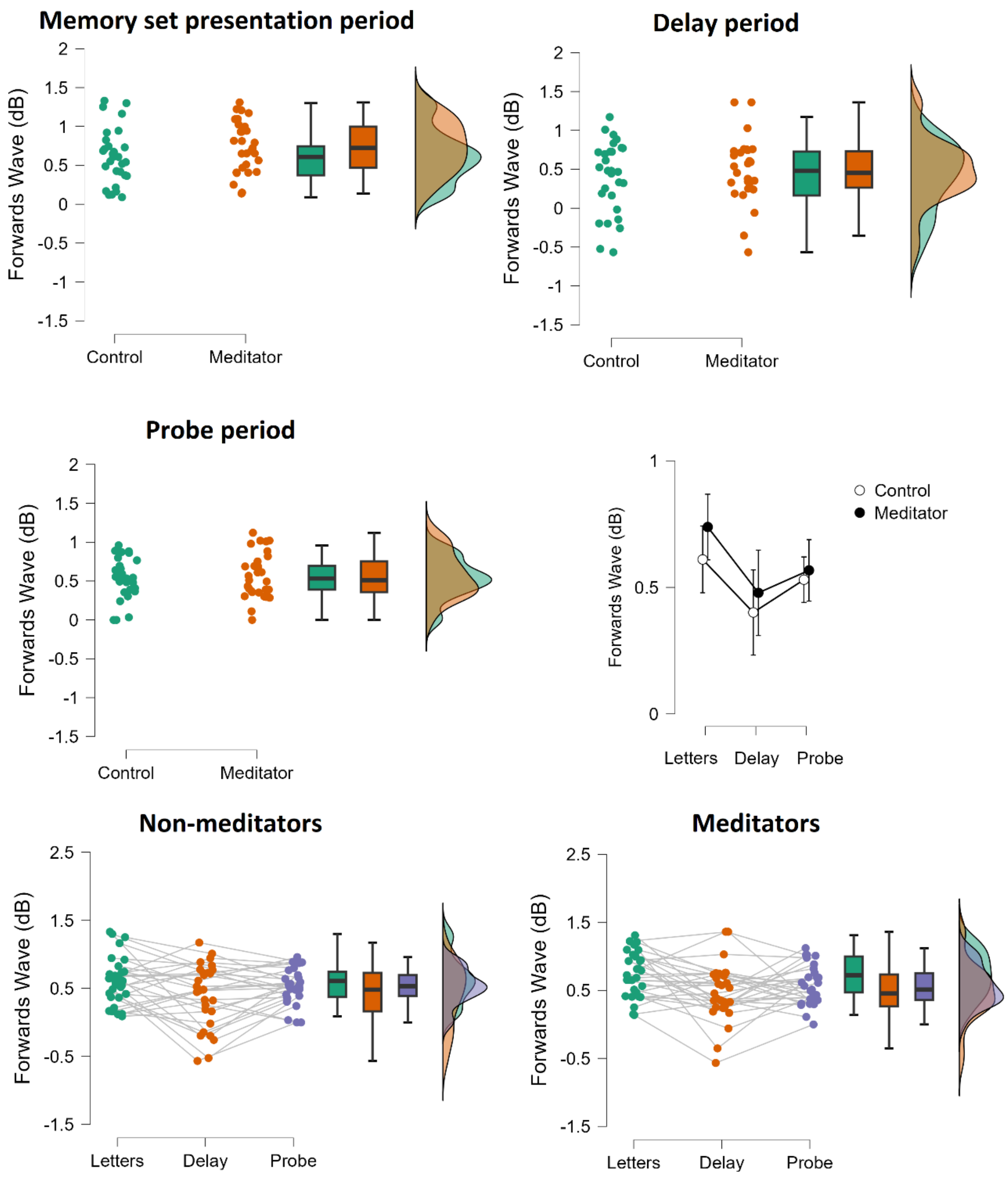

Figure S1. Forwards wave strength (decibels – dB) from the Sternberg working memory task for each group, separated by the task period (top and middle), and each task period, separated by group (bottom). Note that despite the same between group pattern as the other datasets was present within the memory set letter presentation period, within this task there were no significant main effects or interactions involving group (all $p > 0.09$).

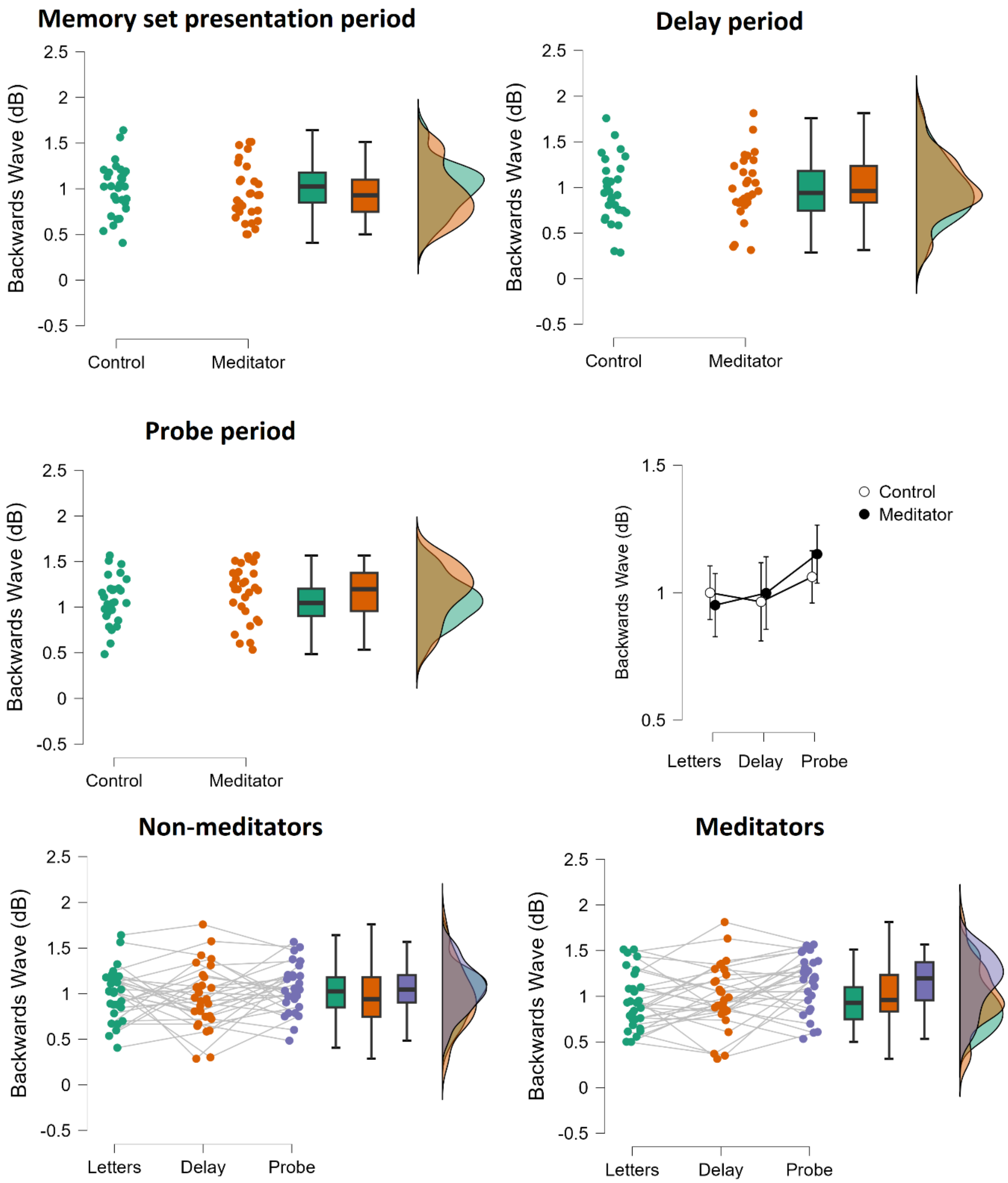

Figure S2. Backwards wave strength (decibels – dB) from the Sternberg working memory task for each group, separated by the task period (top and middle), and each task period, separated by group (bottom). Note that there were no significant main effects or interactions involving group (all $p > 0.09$).

It is worth noting that the meditation group showed the same (non-significant) pattern of stronger forwards waves compared to the non-meditator group in all periods of the memory task, and that this pattern towards stronger forwards waves was the largest during the memory set presentation period (in alignment with task demands and the pattern in the within participants analysis factor). It is also worth noting that during the probe presentation period, the non-significant pattern was in the direction of meditators showing stronger backwards waves than non-meditators (again in line with task demands and the pattern in the within participants analysis factor). To assess the potential that the null result might be a product of a smaller sample size, we performed independent samples t-tests of forwards wave strength between groups within the memory set presentation period, and backwards wave strength within the probe presentation period. These tests showed no significant differences, and inconsequential Bayesian evidence for either the null or alternative hypothesis: forwards waves in the memory set presentation period: $t(56) = 1.420$, $p = 0.161$, Cohen's $d = 0.373$, $BF10 = 0.615$, backwards waves in the probe presentation period: $t(56) = 1.190$, $p = 0.239$, Cohen's $d = 0.312$, $BF10 = 0.480$ see Supplementary Materials Figure 1). Given the Bayesian analysis revealed similar evidential support for the alternative and null models, we interpret this finding as not providing an answer to the question of whether meditators show altered travelling wave strengths in the working memory task, with a larger sample size required.

To assess the potential that motor responses influenced the results for the probe period in the working memory task, we conducted an exploratory comparison of potential interactions between forwards and backwards wave strength and Go and No-go trials from the Go/No-go task. Since Go trials required a button press response, while No-go trials required the lack of a motor response, this analysis provides information about whether travelling waves were affected by motor responses. As such, the analysis provided information that enabled us to assess whether button press responses may have influenced results in the probe condition of the Sternberg task. Within this analysis, there was no significant interaction between the Go/No-go condition and the direction of travelling waves: $F(1, 124) = 0.079$, $p = 0.779$, $\eta p^2 < 0.001$, $\eta G^2 < 0.001$. Nor was there a significant interaction between the Go/No-go condition, the direction of travelling waves, and the meditator or non-meditator group: $F(1, 124) = 0.060$, $p = 0.807$, $\eta p^2 < 0.001$, $\eta G^2 < 0.001$. This lack of difference between motor response and non-response conditions in the Go/No-go task suggests that the difference in travelling wave strength between the different periods of the Sternberg working memory task is unlikely to be driven solely by the presence of motor responses in the probe presentation period. As such, we suggest the results from our analysis of the Sternberg working memory task indicate different levels of bottom-up and top-down predictive processing dependent on task demands, in alignment with previous research (Mohan et al., 2024).
